# Contact- and diffusion-based allelopathic interaction of a seaweed with coral holobionts

**DOI:** 10.64898/2026.07.20.739502

**Authors:** Pei Yi Peggy Tang, Joao P. A. Pereyra, Li Keat Lee, Diane Mcdougald, Scott A. Rice, Lindsey K. Deignan, Rebecca J. Case

## Abstract

Shifts from coral-dominated to macroalgal-dominated reef systems have become increasingly common in many coastal regions worldwide. Coral-macroalgal interactions have been shown to be generally detrimental to coral health, with macroalgal allelopathic compounds able to exert serious and even lethal effects on coral at various stages of growth and development. Previous studies have shown that the coral-associated microbial communities play important roles in coral health and mitigating external environmental stress, including macroalgal contact stress. However, it remains unclear whether changes in the coral microbiome have an influence on the Symbiodiniaceae community composition, and if such changes subsequently affect coral health. In this study, we examined changes in both the coral microbiome and the Symbiodiniaceae communities of two Singaporean coral species (*Pocillopora acuta* and *Merulina ampliata*) when exposed to both direct and water-mediated macroalgal contact with *Lobophora* sp. This was investigated using 16S rRNA gene amplicon sequencing to characterize the coral microbiome, and ITS2 variable region sequencing to profile the Symbiodiniaceae communities. Although no significant differences were observed for the coral microbiomes and Symbiodiniaceae communities at both alpha-and beta- diversity levels between control and macroalgal contact treated fragments within each coral species, inter-colony variations in responses to macroalgal contact were observed for both the coral microbiome and Symbiodiniaceae communities of *M. ampliata*. These results suggest that coral colonies vary in the mechanisms that allow mitigation of the effects of macroalgal contact, and in their resilience to macroalgal-induced stress.

## Introduction

Coral and marine algae are two of the major competitors in the marine benthic environment. Their interactions are some of the most-commonly studied (Barott and Rohwer, 2012) as they have been associated with higher instances of coral disease (Nugues *et al*., 2004).

Coral reef systems worldwide have been on a continuous decline for the past few decades, due to a combination of anthropogenic factors and ocean warming (Hoegh-Guldberg *et al*., 2018; Bellwood *et al*., 2019). Recent assessments carried out by the Global Coral Reef Monitoring Network (GCRMN) have reported increased macroalgal cover and concurrent decreases in coral cover in many coastal regions worldwide (Souter *et al*., 2021), thus necessitating more in-depth studies on coral-macroalgal interactions in the current context of climate change and rapid urbanization of reef systems.

Macroalgae exert their effects on corals through shading, direct overgrowth, and allelopathy (Jompa and McCook, 2003). Macroalgal allelopathy (production of chemical compounds that affect the growth and development of other surrounding organisms; Harlin and Rice, 1987; Semmouri et al., 2024) has been shown to produce severe physiological stress and even mortality on corals (Rasher and Hay, 2010). Macroalgal allelopathy has been observed in all three major groups of macroalgae (Green, Brown, and Red algae); however, certain macroalgal species, for example, green algae such as *Chlorodesmis fastigiata* and *Halimeda sp.*, brown algae such as *Lobophora sp*. and *Dictyota sp*., and red algae such as *Galaxaura filamentosa*, have been found to affect a broad range of coral species (Budzałek *et al*., 2021).

Allelopathic compounds have been isolated from macroalgae in past studies, and these include terpenoids from *Galaxaura filamentosa* and *Chlorodesmis fastigiata* (Rasher *et al*., 2011), neurymenolides from *Phaecelocarpus neurymenioides* (Andras *et al*., 2012) and lobophorenols from *Lobophora rosasea* (Vieira *et al*., 2016). Many identified allelopathic compounds are lipophilic and are transmitted from the macroalgal surfaces to the coral surfaces via direct contact (Longo and Hay, 2017). These secondary metabolites exhibit diverse bioactivities, including antibacterial (e.g. Tolpeznikaite *et al*., 2021), antifungal (e.g. Shobier *et al*., 2016), antiviral (e.g. Castillo *et al*., 2020), antiprotozoal (e.g. Cantillo-Ciau *et al*., 2010) and cytotoxic (e.g. Hamann and Scheuer, 1993) properties (e.g. Dahms and Dobretsov, 2017; Biris-Dorhoi *et al*., 2020).

Past studies have shown that direct macroalgal contact exerts negative effects on coral health and growth (Nugues *et al*., 2004; Rasher and Hay, 2010; Vega Thurber *et al*., 2012) and inflict damage on the coral, causing shifts towards pathogenic microbial communities in the coral microbiome (Morrow *et al*., 2013; Morrow *et al*., 2017). However, there is evidence that water-mediated allelopathy of macroalgal water-soluble compounds may also affect the coral microbiome. Morrow *et al*. (2013) reported microbiome changes even when corals and macroalgae were separated by 5 cm of seawater, possibly due to waterborne compounds mediating these interactions. Smith *et al*. (2006) suggested the production and release of dissolved organic carbon (DOC) into the surrounding water column as an indirect way in which macroalgae could affect coral health through hypoxia, explained by the DDAM model (DOC, Disease, Algae and Microorganism model; Haas *et al*., 2016). Although hydrophilic extracts from macroalgae have varying degrees of antimicrobial and growth-enhancing activities in bacterial assays (Morrow *et al*., 2011), it remains unclear whether these compounds influence coral-associated microbial communities *in situ*. Furthermore, for coral-macroalgal interactions, most studies have been largely focused on changes in the coral microbiome during macroalgal contact (Vega Thurber *et al.,* 2012; Morrow *et al*., 2013; Fong *et al*., 2020). Little is known about possible changes in the Symbiodiniaceae communities during coral-macroalgal contact.

Here, we characterized changes in the coral microbiome and Symbiodiniaceae communities of *Merulina ampliata* and *Pocillopora acuta*, both collected from two different locations within an urbanized coral reef system, in response to different forms of macroalgal contact. To do so, both 16S rRNA gene sequencing and ITS2 amplicon sequencing was used to characterize the responses in the coral microbiomes and Symbiodiniaceae communities during direct and water-mediated macroalgal contact. Consequently, this study clarifies how macroalgal stress disrupts coral holobiont stability and affects the health of the coral host.

## Results

### Coral microbiomes differ by species and across time

Corals of two different species were sourced from two different locations for experiments in aquaria (Figures 1 and 2). To look at their microbiomes, DNA was extracted from samples of their tissue at different time points for amplification and sequencing of partial 16S genes (amplicons). Amplicon sequence variants (ASVs) from samples were then analyzed using PERMANOVA. This analysis showed that *P. acuta* and *M. ampliata* microbiomes differed significantly (*F*(3,169) = 24.5, *p* < 0.05). Although the coral microbiomes were not significantly different based on their sampling locations alone (*P. acuta*: *F*(1,48) = 0.994, *p* > 0.05; *M. ampliata*: *F*(1,48) = 1.17, *p* > 0.05), the coral microbiomes differed significantly across experimental time points as shown in Figure 3 (*P. acuta*: Pulau Satumu: *F*(4,20) = 1.71, *p* < 0.05, Kusu Island: *F*(4,20) = 1.80, *p* < 0.05; *M. ampliata*: Pulau Satumu: *F*(4,20) = 1.51, *p* < 0.05, Kusu Island: *F*(4,20) = 1.91, *p* < 0.05). Because of the highly dynamic nature of coral microbiomes across time, comparisons were made between the control and treated microbiomes taken at the same experimental time point. There were no significant differences between the non-treated control, the water-mediated contact and the direct contact microbiomes at the alpha-diversity level (species richness and evenness) (Tables S1 and S2) and at the beta-diversity level (Table 1) for both *P. acuta* and *M. ampliata*. The only significant differences found were between field samples / samples in aquaria before the experiment was conducted and microbiomes during the experiments (water-mediated contact and direct contact) (Table S2).

**Figure 1.**
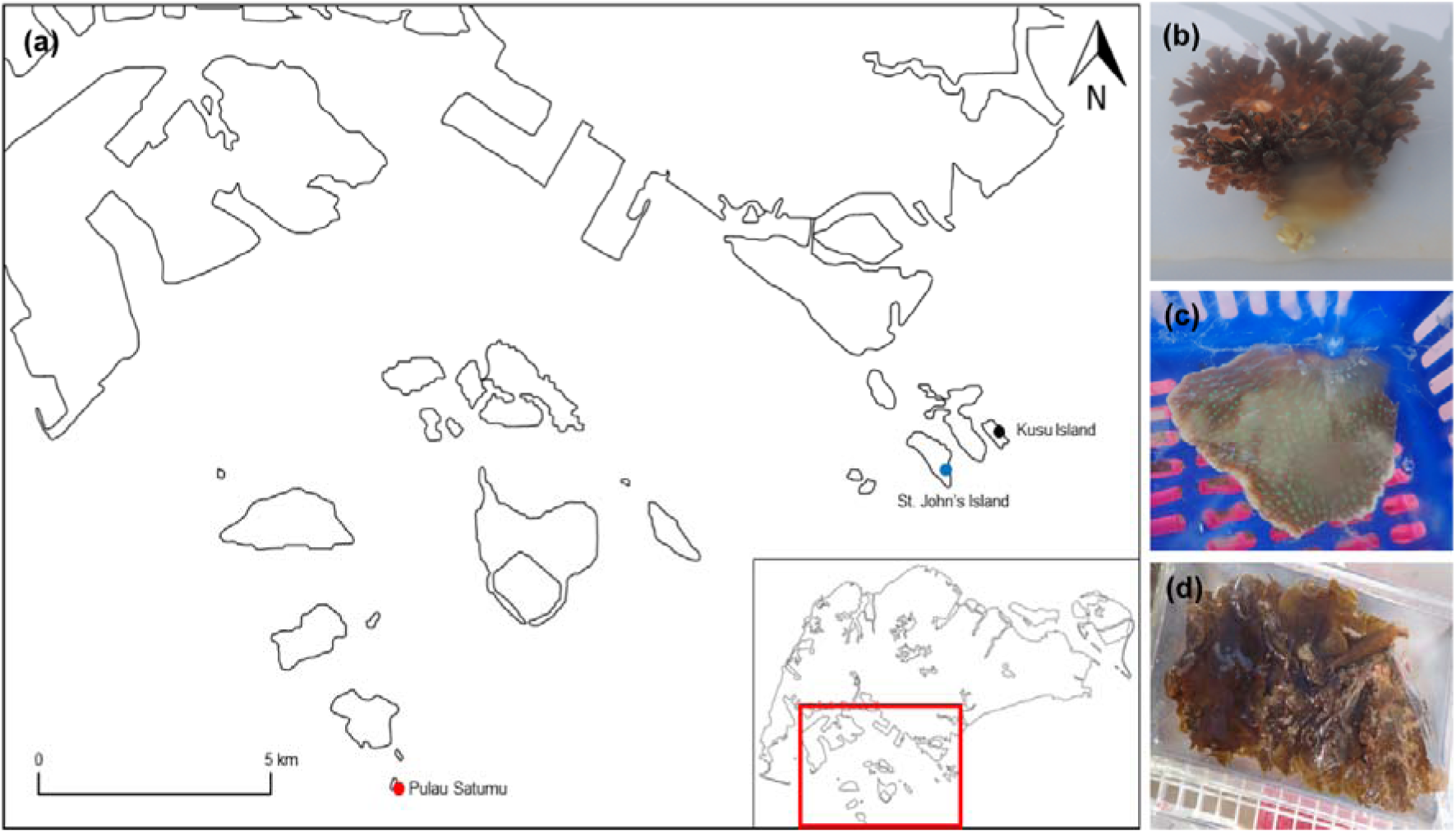
Sampling locations and the coral and macroalgal species used in the study. (a) Both sampling sites Pulau Satumu (red dot) and Kusu Island (black dot), in Singapore, are approximately 15.1 km apart. The St John’s Island National Marine Laboratory (SJINML) where the mesocosm experiments took place, is located on St John’s Island (blue dot). (b) *Pocillopora acuta*, (c) *Merulina ampliata* and, (d) *Lobophora sp*.

**Figure 2.**
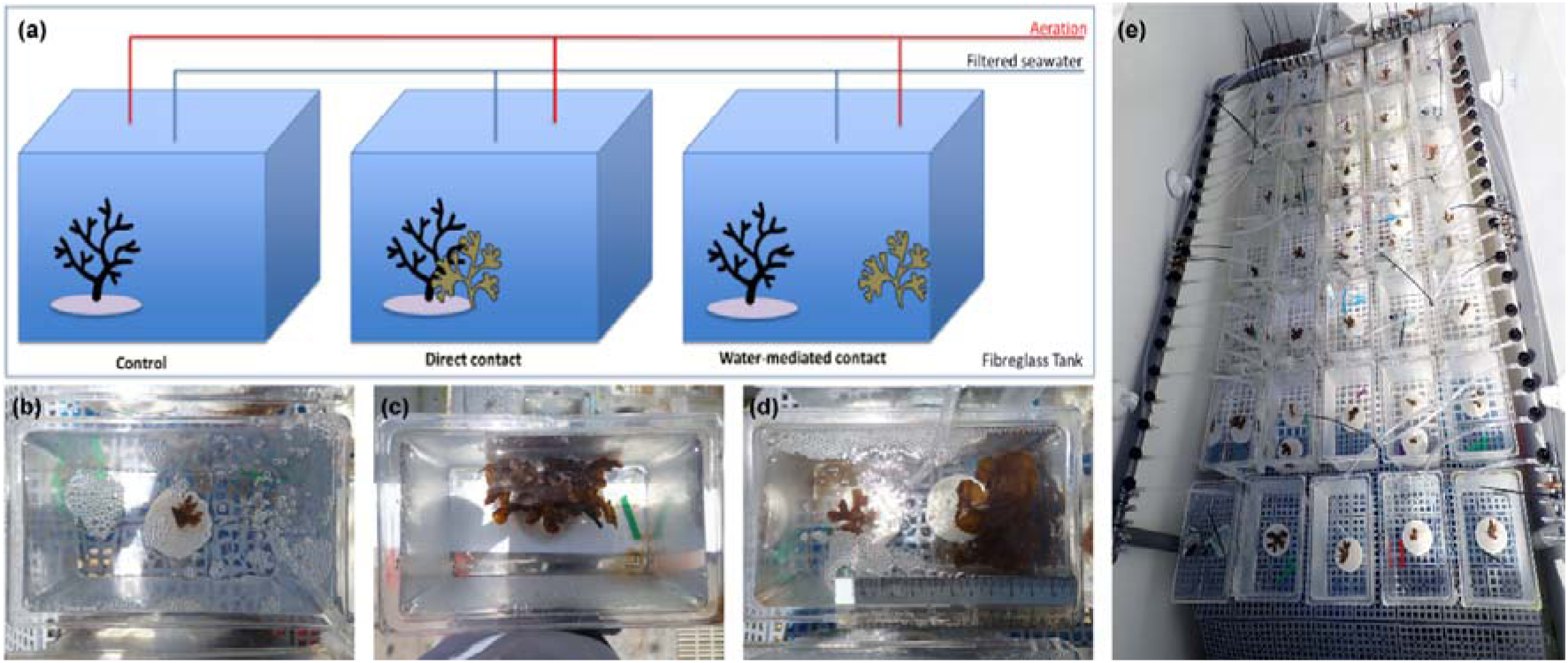
Experimental setup of the mesocosm system in which we manipulate the seaweed-coral contact-dependent and -independent allelopathic interaction. (a) Not-to-scale schematic diagram of the mesocosm setup in the fiberglass holding tank with the different macroalgal contact treatments. The corresponding actual treatment tank photos are below, showing the (b) non-treated control, (c) direct contact-treated and (d) water-mediated contact-treated tanks. Panel (e) shows the actual arrangement of the treatment tanks in the fiberglass holding tank.

**Figure 3.**
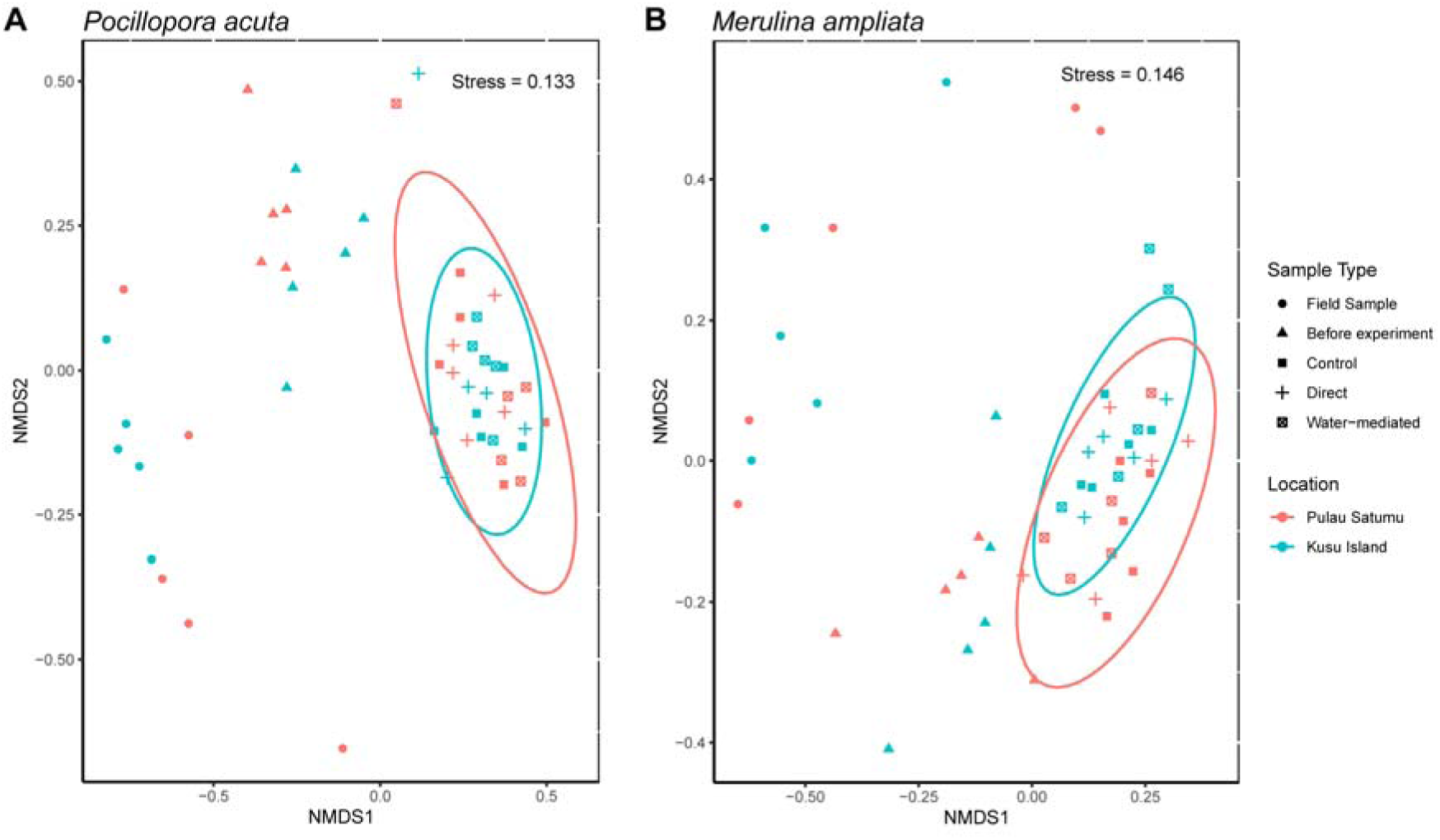
The coral microbiomes of two species at two locations, and not significantly impacted by the coral-seaweed allelopathic interaction. Both (a) *P. acuta* and (b) *M. ampliata* microbiomes shifted from the initial sampling time point, to the end of the experiment. However, the experimental microbiomes (non-treated control, direct contact and water-mediated contact) were more similar, despite differences in sampling locations.

**Table 1.** Summary of PERMANOVA and PERMDISP analyses (with colony strata) for *P. acuta* and *M.ampliata* mesocosm experiments between non-treated control and treated microbiomes based on location.

| Species | Location | PERMANOVA |  | PERMDISP |  |
| --- | --- | --- | --- | --- | --- |
|  |  | *F* | *P<sub>r</sub>* (> *F*) | *F* | *P<sub>r</sub>* (> *F*) |
| *P. acuta* | Pulau Satumu | 1.00 | 0.24 | 0.13 | 0.94 |
|  | Kusu Island | 0.95 | 0.37 | 1.53 | 0.26 |
| *M. ampliata* | Pulau Satumu | 0.92 | 0.69 | 0.69 | 0.53 |
|  | Kusu Island | 0.97 | 0.70 | 2.33 | 0.14 |

\* Bold values indicate significant differences when $p < 0.05$ .

### Coral microbiome changes during macroalgal contact

Although there was no significant change in the coral microbiome after macroalgal contact at the community (alpha diversity) level, direct contact visibly affected the coral, with lighter patches at points of contact (Figure 4).For both *P. acuta* and *M. ampliata*, the dominant phyla were Proteobacteria and Bacteroidota, followed by Verrucomicrobiota and Planctomycetota, with other bacterial phylum including Desulfobacterota, Firmicutes, Myxococcota and other taxa showing relative abundances less than 1% of the total read count (Figure 5). Variations for specific phyla between the non-treated control and treated microbiomes across different coral colonies originating from a specific sampling site could also be observed for both *P. acuta* and *M.ampliata* (Figure 5). These variations, however, were not consistent across replicate colonies of the same corals and the specific phylum affected. For example, the three most notable changes observed after direct macroalgal contact are: 1) an increase in Campilobacterota is seen for one of the five Kusu *P. acuta* colonies (Kusu3, Figure 5B); 2) an increase in Desulfobacterota and Firmicutes in one of five Kusu *M. ampliata* colonies (Kusu3, Figure 5B); 3) an increase in Cyanobacteria in one of five Satumu *M. ampliata* colonies (Satumu5, Figure 5C).

**Figure 4.**
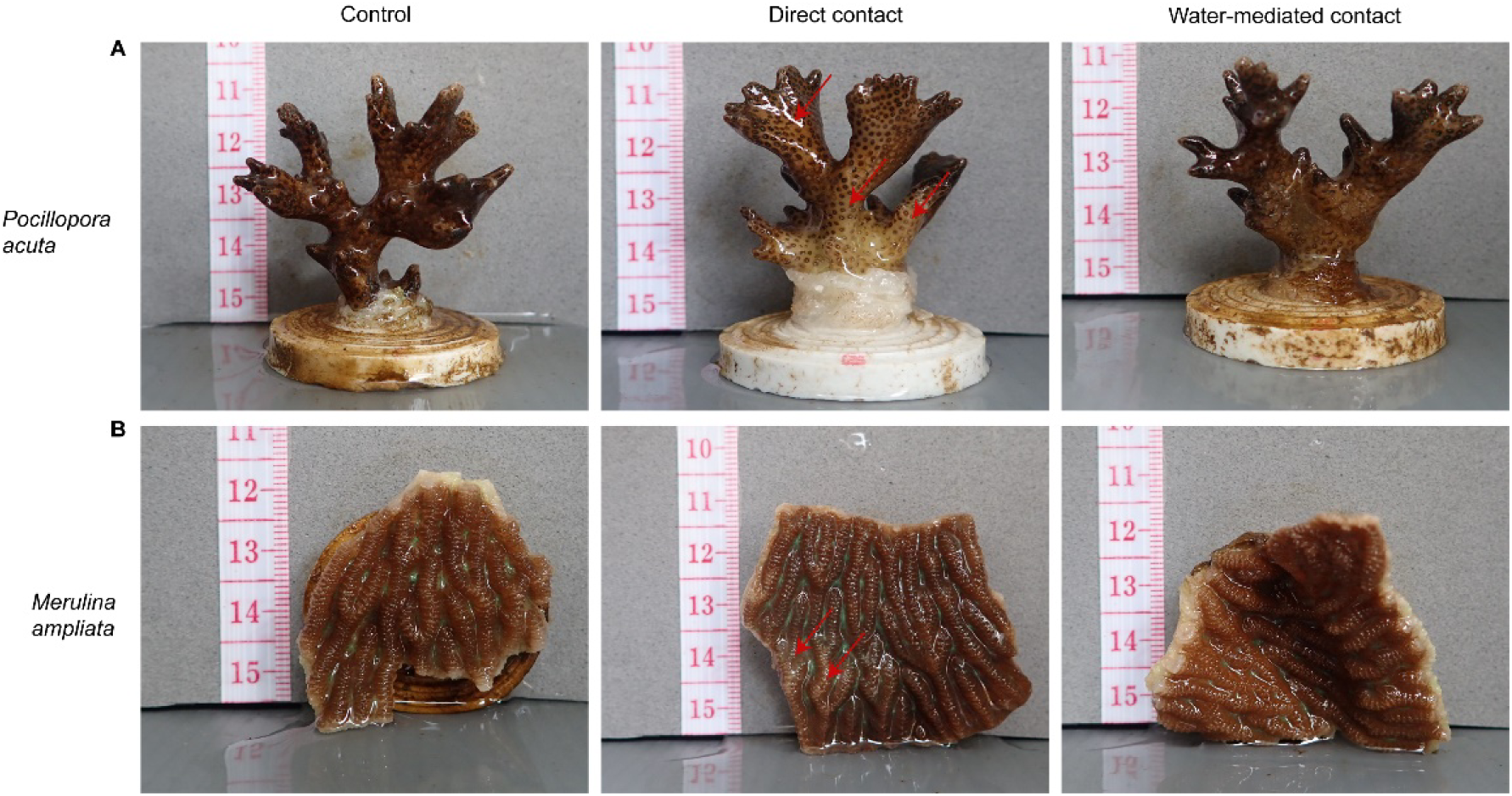
Direct contact between seaweed-coral produces bleaching, while diffusion-mediated allelopathy does not produce coral bleaching. Top: (a) *Pocillopora acuta*, (b) *Merulina ampliata*. The red arrows in the photos of the coral fragments exposed to direct macroalgal contact for both coral species indicate significantly lighter areas where the macroalgae was in direct contact with the coral surface, as compared to the non-treated control and water-mediated macroalgal contact fragments.

**Figure 5.**
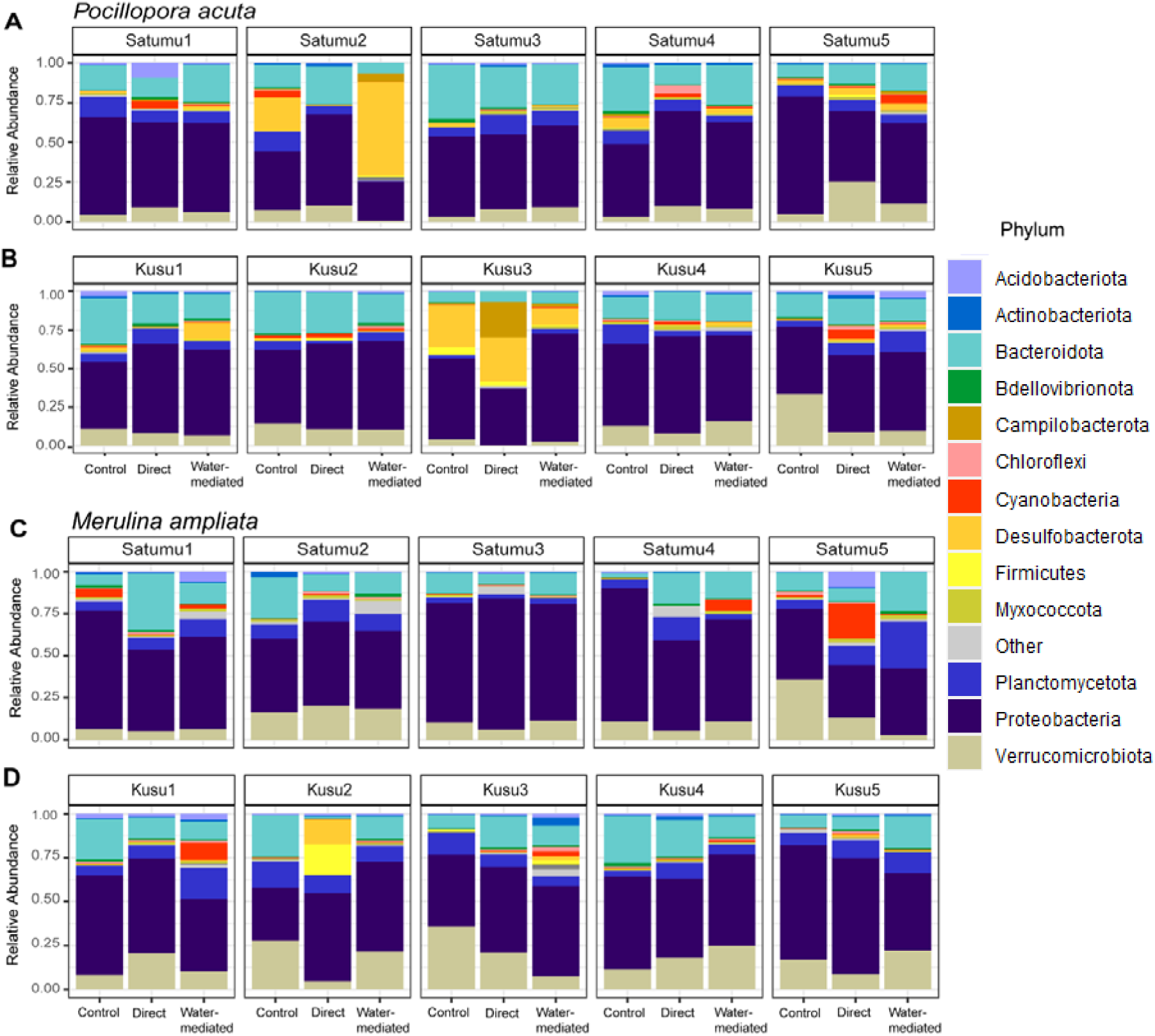
Phylum-level changes in the coral microbiome during macroalgal contact across individual colonies and sampling location for *P. acuta* and *M. ampliata.* From top: *P. acuta*: (a) Pulau Satumu, (b) Kusu Island, *M. ampliata*: (c) Pulau Satumu, (d) Kusu Island. Satumu1-Satumu5 and Kusu1-Kusu5 refer to a fragment from each of the five individual corals sampled from Pulau Satumu and Kusu Island, respectively. Variation of changes in phyla in response to macroalgal contact was observed across both coral species, sampling locations and type of macroalgal contact

At the individual ASV level, several ASVs differed between control and treatment (Table S3). In both coral species, ASVs associated with the phyla Proteobacteria, Bacteroidota, Verrucomicrobiota and Planctomycetota accounted for the majority of the observed changes. Additional changes were detected in ASVs assigned to the phyla Desulfobacterota, Bdellovibrionota, Cyanobacteria, Acidobacterota, Chloroflexi, Actinobacteria, Campilobacterota, Myxococcota, Firmicutes, Deinococcota, Nitrospirota, Dependentiae and Fusobacterota (Table S3).

At a finer taxonomic resolution for *P. acuta*, relative abundances of ASVs assigned to *Desulfobacter sp.*, Candidatus Actinomarina (phylum Actinobacteriota), the family Alteromonadaceae and the family Roseobacteraceae were increased in microbiomes exposed to both macroalgal contact types (Table S3). In colonies from Pulau Satumu, direct macroalgal contact was associated with increased relative abundances of ASVs assigned to *Erythrobacter sp*., *Zeaxanthinibacter sp*., *Woeseia sp*., *Lentimonas sp.*, and *Aquibacter sp*. Water-mediated macroalgal contact was associated with an increase in ASVs assigned to *Synechococcus sp*. For both types of macroalgal contact, relative abundances of ASVs associated with *Muricauda sp.* and *Maricurvus sp*. decreased. In *P. acuta* colonies from Kusu Island, ASVs assigned to *Pseudomonas sp*., *Lentimonas sp*., Candidatus Endobugula (family Cellvibrionaceae), Candidatus Actinomarina (phylum Actinobacteriota), and the family Roseobacteraceae, increased in relative abundance under both macroalgal contact treatments. Direct macroalgal contact additionally resulted in increased relative abundances of ASVs assigned to *Kordiimonas sp*., *Desulfobacter sp*., *Cyanobium sp*., *Aestuariibacter sp*. and the family Arcobacteraceae (phylum Campilobacterota). Water-mediated macroalgal was associated with increased relative abundances of ASVs assigned to *Ruegeria sp.*, *Erythrobacter sp*., *Woeseia sp.* and the family Alteromonadaceae. For both types of macroalgal contact, relative abundances of ASVs associated with *Muricauda sp*., *Arenicella sp*. and the family Vibrionaceae decreased (Table S3).

In *M. ampliata*, several ASVs also exhibited changes in relative abundances in response to macroalgal contact (Table S4). In colonies from Pulau Satumu, ASVs assigned to *Pseudoalteromonas sp*., *Arenicella sp*., *Aliikangiella sp*., *Maricurvus sp.* and the family Roseobacteraceae increased in relative abundance under both macroalgal contact types. Direct macroalgal contact additionally resulted in increased relative abundances of ASVs assigned to the OM27 clade (phyla Bdellovibrionota), the family Vibrionaceae and *Ruegeria sp.* Water-mediated macroalgal contact was associated with increased relative abundances of ASVs assigned to *Woeseia sp*., *Lentimonas sp.*, *Labrenzia sp*., *Synechococcus sp*. and Candidatus Endobugula (family Cellvibrionaceae). Contrasting, ASVs assigned to *Trichodesmium sp.* decreased in relative abundance under both types of macroalgal contact. In *M. ampliata* colonies from Kusu Island, ASVs assigned to *Lentimonas sp*. and *Oleiphilus sp*. increased in relative abundance under both macroalgal contact treatments. Direct macroalgal contact additionally resulted in increased relative abundances of ASVs assigned to *Leisingera sp*. and *Erythrobacter sp*. (Table S4). Water-mediated macroalgal contact was associated with increased relative abundances of ASVs assigned to *Altererythrobacter sp*., *Muricauda sp*., *Woeseia sp*., *Marinobacterium sp.*, the family Flavobacteriaceae and the family Roseobacteraceae. In contrast, ASVs assigned to the order Chlamydiales (family Simkaniaceae) and Candidatus Endobugula (family Cellvibrionaceae) decreased in relative abundance under both types of macroalgal contact.

### Effects of macroalgal contact on the coral Symbiodiniaceae communities

Using Symportal (Hume *et al*., 2019), we determined that the dominant Symbiodiniaceae taxon present in *P. acuta* belonged to the genus *Durusdinium sp*. (clade D), specifically, *Durusdinium glynii* (LaJeunesse *et al*., 2018) (Figure 6). Low-abundance *Cladocopium sp*. (clade C) ITS2 profiles were found only in colony Satumu5 (non-treated control) (Figure 6). In *M. ampliata*, *Cladocopium sp*. (clade C) was the dominant genus, with *Durusdinium sp*. (clade D) also detected in the colonies Kusu2 (all contact types), Kusu4 (non-treated control) and Kusu5 (all contact types), with a low-abundance population in Satumu2 (non-treated control) (Figure 6). In both coral species, the detected *Cladocopium* ITS2 types (indicative of distinct subclades) included C3, C3, C115 and C21ab, whereas the detected *Durusdinium* ITS2 types included D1, D1cn, D1co, D1cp, D1cr, D1n and D2d (Figure 6). In *M. ampliata*, although the dominant *Durusdinium* ITS2 profiles (D1/D2d) was always present, there were variations in the composition of the low-abundance ITS2 types across different individual colonies (Figure 6).

**Figure 6.**
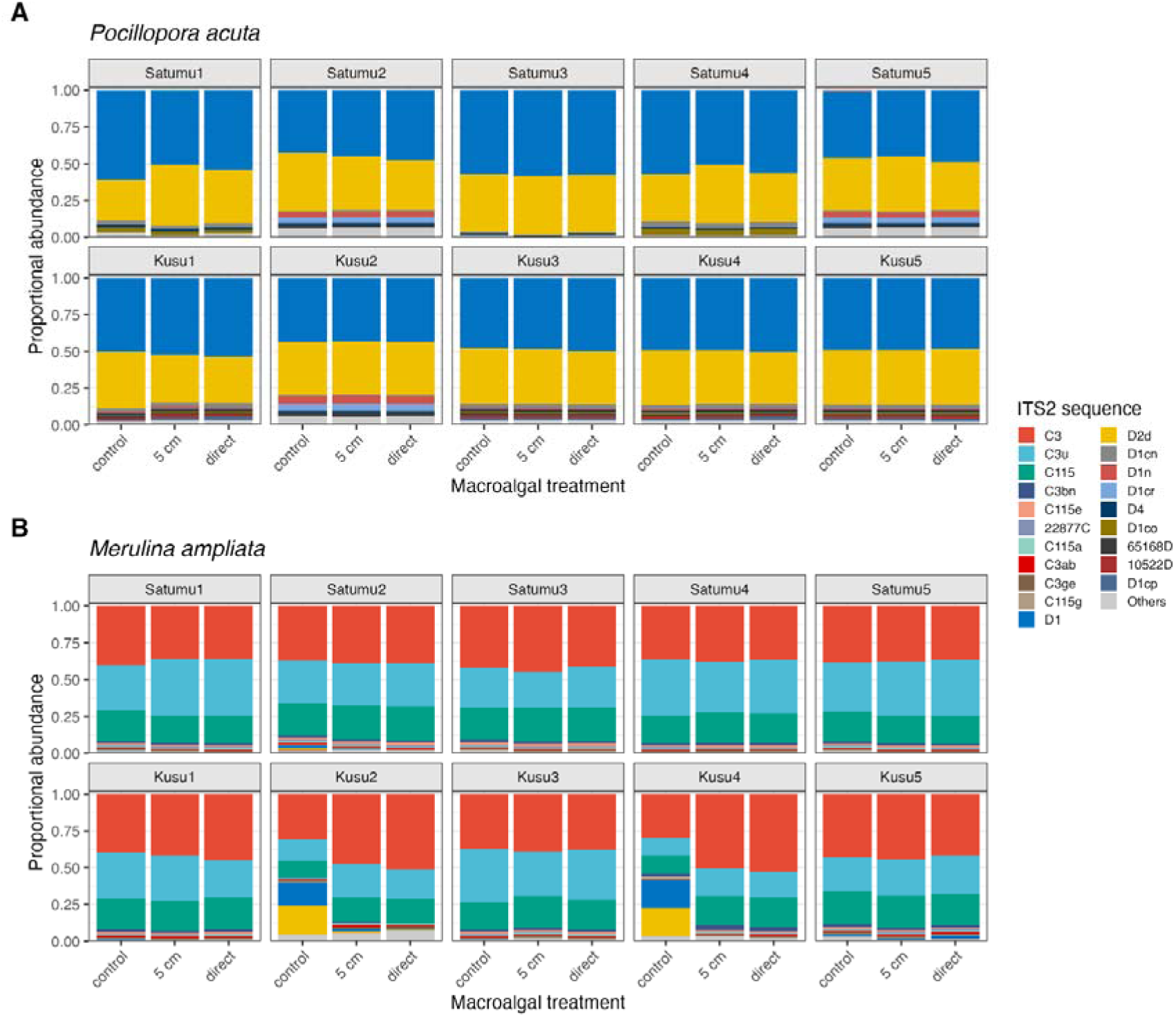
Characterization of the Symbiodiniaceae communities in both *P. acuta* and *M. ampliata* during macroalgal contact. From top: (a) *P. acuta* Pulau Satumu and Kusu Island, (b) *M. ampliata* Pulau Satumu and Kusu Island. Satumu1-Satumu5 and Kusu1-Kusu5 refer to a fragment from each of the five individual corals sampled from Pulau Satumu and Kusu Island, respectively. Distribution of the Symbiodiniaceae clades is coral species-specific and may vary across individual colonies within a particular location. Determination of Symbiodiniaceae subclades was done using Symportal (Hume *et al*., 2019).

Significant differences in Symbiodiniaceae communities (Chao1 index) were observed among the Pulau Satumu *P. acuta* treatment (Table S5). Post-hoc Tukey HSD tests showed that the observed differences were due to differences between Symbiodiniaceae communities exposed to water-mediated macroalgal contact and direct macroalgal contact, and between Symbiodiniaceae communities exposed to water-mediated macroalgal contact and those from field samples (Table S6). No significant differences were observed for the other alpha diversity indices from either Pulau Satumu or Kusu Island (Table S5). In *M. ampliata*, no significant differences in Chao1, Shannon-Wiener or Inverse Simpson indices were observed among the macroalgal contact treatments at either Pulau Satumu or Kusu Island (Table S5). PERMANOVA analyses showed that differences in Symbiodiniaceae communities in both coral species were colony-dependent, and not affected by the type of macroalgal contact present (Table 2).

**Table 2.** Summary of PERMANOVA and PERMDISP analyses for *P. acuta* and *M.ampliata* Symbiodiniaceae communities between non-treated control and treated samples based on location.

| Species | Location | Variable | PERMANOVA |  | PERMDISP |  |
| --- | --- | --- | --- | --- | --- | --- |
|  |  |  | *F* | *P<sub>r</sub>* (> *F*) | *F* | *P<sub>r</sub>* (> *F*) |
| *P. acuta* | Pulau Satumu | Colony (One-way) | 6.3898 | **0.001** | 0.175 | 0.944 |
|  |  | Macroalgal Contact Type (with colony strata) | 0.5113 | 0.401 | 0.8747 | 0.484 |
|  | Kusu Island | Colony (One-way) | 3.2251 | **0.003** | 0.4598 | 0.767 |
|  |  | Macroalgal Contact Type (with colony strata) | 2.04 | 0.06725 | 1.898 | 0.134 |
| *M. ampliata* | Pulau Satumu | Colony (One-way) | 2.4863 | **0.005** | 1.1084 | 0.395 |
|  |  | Macroalgal Contact Type (with colony strata) | 1.4288 | 0.053 | 0.9196 | 0.476 |
|  | Kusu Island | Colony (One-way) | 4.0779 | **0.001** | 1.0301 | 0.428 |
|  |  | Macroalgal Contact Type (with colony strata) | 0.9141 | 0.133 | 0.6713 | 0.621 |

\* Bold values indicate significant differences when $p < 0.05$ .

## Discussion

### Macroalgal contact has minimal effect on the coral microbiome at both alpha- and beta-diversity levels

In this study, neither direct nor water-mediated macroalgal interactions resulted in significant changes in the overall alpha or beta diversity of the microbiomes associated with *P. acuta* and *M. ampliata*. However, changes in the relative abundances of specific bacterial ASVs were detected, suggesting that macroalgal contact elicited localized shifts within the microbiomes rather than large-scale changes. Previous studies have reported increased species richness and reduced community evenness after macroalgal contact (Zaneveld *et al*., 2016; McDevitt-Irwin *et al.,* 2017; Fong *et al*., 2020), attributed to a reduced capacity of the coral host to maintain the integrity of the coral microbiome (McDewitt-Irwin *et a*l., 2017). The destabilization of the coral microbiome also leads to increased dispersion among microbiome communities following disturbance (Zaneveld *et al*., 2016). However, the absence of community-level responses in our study suggests that the microbiomes of *P. acuta* and *M. ampliata* remained relatively stable following both types of macroalgal contact, which could be attributed to the high stress levels in the urban reef environment from which the corals originated.

Despite the lack of significant community-level changes in the coral microbiomes from both coral species during macroalgal contact, minor localized tissue bleaching was observed for direct contact-treated fragments, consistent with observations from Fong *et al*. (2020). This suggests that physiological responses may occur even in absence of major changes of the associated microbial communities. Given that pulse-amplitude modulated (PAM) fluorometry is widely used as an indicator of coral health (Ralph *et al*., 1999; Fitt *et al*., 2001), incorporating PAM measurements in future studies would facilitate changes in microbial community composition to be more directly related to physiological changes in corals.

### Changes in relative abundances of ASVs associated with specific bacterial groups were observed during macroalgal contact

Although no significant change in the microbial community structure was observed at the alpha- and beta-diversity levels, changes in the relative abundance of specific groups of bacterial ASVs were still observed for both coral species. Many of these ASVs were assigned to members of the phyla Proteobacteria, Bacteroidota, Planctomycetota and Verrucomicrobiota. Some of these bacterial groups have been associated with nutrient cycling and nutrition, for example, members of the family Cellvibrionaceae (Spring *et al*., 2015) and *Lewinella sp*. (McIlroy and Nielsen, 2014) have been implicated with complex carbohydrate hydrolysis, with members of the former capable of nitrogen fixation (Suarez *et al*., 2014) and identified as important endosymbionts in other organisms such as shipworms (Brito *et al*., 2018; Lucena *et al*., 2020). *Lentimonas sp*. are known to degrade algal fucoidan (Sichert *et al*., 2020). Other taxa, including *Synechococcus*, members of the family Kiloniellaceae and *Woeseia* have been reported to be involved in nitrogen fixation (Lesser *et al*., 2007), denitrification (Imhoff and Wiese, 2014), and sulfur oxidation (Muβmann *et al*., 2017), respectively. Some of these bacterial groups may also be associated with protective functions. Members of the family Kiloniellaceae have been reported to produce antimicrobials (Imhoff and Wiese, 2014), while some members of *Erythrobacter sp.* (Setiyono *et al*., 2019) and *Muricauda sp.* (Motone *et al*., 2020) produce carotenoids that may provide protection against heat and light stress. Likewise, *Ruegeria* species have been shown to inhibit pathogenic *Vibrio* spp. (Miura *et a*l., 2019). Additionally, members of the families Micavibrionaceae (Mookherjee and Jurkevitch, 2022) and Bdellovibrionaceae (Martin, 2022) are predatory bacteria that prey on Gram-negative bacteria, potentially altering the coral microbial community composition (Mookherjee and Jurkevitch, 2022).The observed shifts of these ASVs suggests that macroalgal contact may selectively influence bacterial taxa with reported roles in nutrient cycling and host protection, and may contribute to maintaining microbiome stability despite macroalgal contact stress.

In addition to potential beneficial taxa, there were also increases in ASVs associated with the families Roseobacteraceae, Campilobacteraceae, *Vibrio sp*., and the sulfate-reducing genus *Desulfobacter sp*. Several of these bacterial groups have been reported to increase in diseased corals or associated with coral disease (Kellogg *et al*., 2013; Pootakham *et al*., 2019; Bourne et al., 2011), with various *Vibrio* species recognized as coral pathogens (Kushmaro *et al*., 1997; Morrow *et al*., 2017). Morrow *et al*. (2017) also showed that exposure to macroalgal extracts can promote shifts in coral microbiomes towards potentially pathogenic taxa. However, it is not known if these bacterial groups found in our study, especially members of the Roseobacteraceae and *Vibrio sp*., are pathogenic or non-pathogenic, as the taxonomic resolution of 16S rRNA gene amplicon sequencing doesn’t allow these groups to be distinguished reliably. Furthermore, some of these bacterial groups, including members of the Roseobacteraceae and *Vibrio sp*., are commonly found in healthy coral microbioes, playing important ecological roles such as sulfur cycling and dimethylsulfoniopropionate (DMSP) metabolism (Raina *et al*., 2009), and hence their increased abundance should not be interpreted as an increase in pathogens. ASVs assigned to the phylum Planctomycetota, include *Blastopirellula sp*., *Rhodopirellula sp*., *Rubripirellula sp*. and *Pirellula sp*. Although little is known about their functions in the coral microbiome, they are known to utilize sulphated polysaccharides produced by macroalgae (Lage and Bondoso, 2014). Their enrichment may therefore be due to increased amounts of macroalgal-derived polysaccharides in the surrounding seawater, although testing this hypothesis would require the integration of macroalgal-derived microbiome datasets with dissolved organic carbon measurements.

We did not detect significant changes in ASVs assigned to *Endozoicomonas sp*. in response to macroalgal contact. *Endozoicomonas sp*. is regarded as an important member of the coral microbiome, due to its frequent association with healthy corals and putative functional roles such as nutrient cycling and host metabolism (Neave *et al*., 2017; Pogoreutz and Ziegler, 2023), as well as their ubiquity in numerous coral species (McCauley *et al*., 2023). Although *Endozoicomonas* was detected in both coral species in our study, they were present at very low relative abundances (< 0.05%). It may be possible that their low abundance was due to environmental stressors present in urban reefs, as reductions in abundance have previously been reported under stressful conditions (Ziegler *et al*., 2017). However, host-*Endozoicomonas* relationships may also differ between coral species, coral host genetic lineage and sampling locations, indicating possible differing host- and site-specific strategies for maintaining stability (Hochart *et al*., 2023).

### Symbiodiniaceae symbionts are not affected by macroalgal contact in P. acuta but can be lost in M. ampliata

The Symbiodiniaceae assemblages of both *M. ampliata* and *P. acuta* were dominated by *Cladocopium* and *Durusdinium* ITS2 type profiles, consistent with previous surveys of corals from Singapore and Malaysia (Tanzil *et al*., 2016; Leveque et al. 2019; Poquita-Du et al. 2020; Smith et al. 2020; Lee et al. 2022; Ong et al. 2022) (Figure 6). *M. ampliata* predominantly hosted the ITS2 type profile C3/C3u-C115, which has also been reported from a wide range of coral genera including *Platygyra, Pectinia, Acropora* and *Montipora,* indicating that this *Cladocopium* ITS2 type profile is broadly distributed across diverse coral hosts (Smith et al. 2020; Lee et al. 2022; Ong et al. 2022). In contrast, *P. acuta* was dominated by *Durusdinium,* with profiles based mainly on D1 and D2d types, again consistent with previous observations in the region (Poquita-Du et al. 2020; Ong et al. 2022). Macroalgae such as *Lobophora* produce allelochemicals that can induce coral bleaching following direct contact, although these effects are species-specific (Rasher and Hay 2010; Rasher et al. 2011; Vieira et al. 2016). In this study, both coral hosts (*P. acuta* and *M. ampliata*) exhibited signs of bleaching and changes in their bacterial microbiomes following contact with *Lobophora.* However, the Symbiodiniaceae communities responded differently between the coral species. In *P. acuta*, the bacterial microbiome shifted whereas the Symbiodiniaceae community remained relatively stable. In contrast, *M. ampliata* exhibited changes in both its bacterial microbiome and Symbiodiniaceae community. *P. acuta* was dominated by *Durusdinium,* a genus widely associated with tolerance to thermal stress and other environmental stressors (Stat et al. 2013; Poquita-Du et al. 2020), which may also confer greater resilience to macroalgal allelochemical exposure than *Cladocopium-*dominated *M. ampliata.* By comparison, although *M. ampliata* hosted co-dominant *Cladocopium* (C3, C3u, C115) and *Durusdinium* (D1, D2d) profiles under control conditions, this mixes assemblage was disrupted following contact with *Lobophora*, suggesting that allelochemical exposure altered the equilibrium of its Symbiodiniaceae community. Notably, although both species exhibited bleaching, only *M. ampliata* showed a compositional shift in its Symbiodiniaceae, suggesting that *Lobophora* allelochemicals may primarily damage coral host tissues, with changes in Symbiodiniaceae composition occurring as a secondary consequence of host damage. This interpretation is consistent with reports that diverse and abundant Symbiodiniaceae, including free-living and in-hospite taxa, can occur on the surfaces of macroalgae (Fujise et al. 2021; Million et al. 2025), indicating that macroalgae and their surface chemistry are not inherently hostile to these dinoflagellates.

## Conclusion

In conclusion, this study demonstrates that short-term direct and water-mediated macroalgal contact resulted in minimal changes to the bacterial microbiome and the Symbiodiniaceae communities of *P. acuta* and *M. ampliata*. However, species-specific differences in both the bacterial microbiomes and Symbiodiniaceae communities were evident, and changes in the relative abundances of specific bacterial ASVs and low-abundance Symbiodiniaceae ITS2 types were detected following macroalgal contact. These findings suggest that coral responses to macroalgal interactions may comprise of targeted shifts in specific microbial taxa while maintaining the overall structure of the coral holobiont. These variations and changes in the microbiomes and Symbiodiniaceae communities may help the corals better adapt to their local environments, and possibly contribute to resilience against environmental stressors, including macroalgal contact. Further studies, including long-term monitoring studies, would be required to better characterize the effects of these variations on coral health and persistence under increasing environmental stress.

## Materials and Methods

### Environmental collection of coral and algae

Sampling was carried out on January 12^th^ 2021, at two locations, Pulau Satumu and Kusu Island, with mesocosm experiments carried out at the St John’s Island National Marine Laboratory (SJINML) (Figure 1). Two common coral species in Singapore (Huang *et al*., 2009; Fong *et al*., 2020), *Merulina ampliata* and *Pocillopora acuta*, were collected for the experiments. *Lobophora sp.* was chosen, as members of this genus have been found to have allelopathic effects against corals (Vieira *et al*., 2016; Fong *et al*., 2019; Fong *et al*., 2020). Five coral colonies were collected at each sampling site, whereas the macroalgae was collected from Kusu Island. Both corals and macroalgae were transported to SJINML for the mesocosm experiments.

### Mesocosm experiment

The mesocosm setup was based on that as described by Fong et al. (2020). In summary, three treatments were used in this study: direct macroalgal contact, water-mediated contact (coral and macroalga separated by 5 cm between their closest edges), and no contact (control without macroalgae) (Figure 2). For each coral species, five replicates were obtained per location, colony and macroalgal contact treatment, resulting in a total number of 300 replicates for the experiment. Each replicate was housed in individual 18 x 11 x 11 cm acrylic tanks supplied with separate filtered and aerated seawater (Figure 2). The tanks were cleaned every four to five days and repositioned within the fiberglass tank to minimise positional effects. Light intensity and temperature were measured using Onset HOBO UA-002-64 Pendant® Temperature/Light 64K Data Loggers (Bourne, Massachusetts, United States). Corals and macroalgae were acclimated separately for two weeks before the start of the mesocosm experiments. Mesocosm experiments for each coral species ran for two weeks, during which coral colour and bleaching were assessed weekly.

### Sample collection and processing

Coral fragments were destructively sampled, preserved immediately in Zymo DNA/RNA Shield solution (Irvine, California, United States), and transported on ice to the Singapore Centre for Life Sciences Engineering (SCELSE) at Nanyang Technological University (NTU) for further processing. Coral tissue was removed from the skeleton in the preservation solution using a WaterPik. DNA was extracted from the coral tissue using Qiagen DNeasy PowerBiofilm Kit (Hilden, Germany) and the Zymo Clean & Concentrator-25 Kit (Irvine, California, United States). DNA quality was assessed with Thermo Fisher Scientific NanoDrop 2000 Spectrometer (Waltham, Massachusetts, United States) and quantified using the Invitrogen Qubit dsDNA High Sensitivity Kit (Thermo Fisher Scientific; Waltham, Massachusetts, United States) for downstream applications.

### 16S rRNA gene and ITS2 variable region amplicon sequencing

For bacterial community analysis, the V4 region of the 16S rRNA gene was amplified with the 515F-806R primer pair (Caporaso *et al*., 2011; Apprill *et al*., 2015; Parada *et al*., 2016). PCR cycling conditions were: 95 ℃ for 5 min, 37 cycles of 94 ℃ for 30 s, 53 ℃ for 40 s, and 72 ℃ for 1 min, with a final extension step at 72 ℃ for 10 min. Each PCR reaction contained 10 μL of Qiagen HotStar Taq Plus Master Mix (Hilden, Germany), 1 μL each of 10 μM forward and reverse primers, 1 μL 200 μM Bovine Serum Albumin (BSA), 4 μL Ambion nuclease-free water (Thermo Fisher Scientific; Waltham, Massachusetts, United States), 1 μL dimethyl sulfoxide (DMSO) and 10 ng of template DNA.

For Symbiodiniaceae community analysis, the ITS2 variable region was amplified using the SYM_VAR_5.8S2-SYM_VAR_REV primer set (Hume *et al*., 2018). PCR conditions comprised of 15 minutes initial denaturation, followed by 35 cycles of 98 ℃ for 2 min, 98 ℃ for 10 sec, 56 ℃ for 30 sec and 72 ℃ for 30 sec, with a final extension at 72 ℃ for 5 min. Each PCR reaction contained 12.5 μL of Qiagen HotStar Taq Plus Master Mix (Hilden, Germany), 1 μL of 10 μM forward and reverse primers, 8.75 μL Ambion nuclease-free water (Thermo Fisher Scientific; Waltham, Massachusetts, United States), 0.75 μL DMSO and 1 μL of 10 ng μL-1 template DNA.

For both amplicons, PCR products were separated on 2% agarose gel, excised and purified using the Invitrogen PureLink Quick Gel Extraction Kit (Thermo Fisher Scientific; Waltham, Massachusetts, United States), quantified using the Invitrogen Qubit dsDNA High Sensitivity Kit (Thermo Fisher Scientific; Waltham, Massachusetts, United States), and assessed using an Agilent BioAnalyzer with D1000 ScreenTape (Santa Clara, California, United States). Purified amplicons were then submitted to the sequencing facility at SCELSE for 300 bp paired-end sequencing on an 200μM Illumina MiSeq platform (San Diego, California, United States).

### Bioinformatic and statistical analyses

For 16S rRNA gene amplicon, the sequences were processed using *dada2* (Callahan *et al*., 2016) in RStudio (RStudio Team, 2015), following the DADA2 Pipeline (v1.16) (https://benjjneb.github.io/dada2/tutorial.html) to generate amplicon sequence variants (ASVs). Reads were quality-filtered (*trimLeft* = 52,52, *truncLen* = 190,190), dereplicated, merged, and screened for chimeric sequences. Taxonomy was assigned against the SILVA 138.1 SSU Ref Nr99 database (https://zenodo.org/record/4587955#.Y5JKUXZBzIU) (Quast *et al*., 2012; Yilmaz *et al*., 2014) with the ‘assignTaxonomy’ and ‘addSpecies’ functions. The phyloseq objects for subsequent data analyses were generated using phyloseq (v1.32.0) (McMurdie and Holmes, 2013). Samples were rarefied to 95% sequencing depth using the ‘rarefy_even_depth’ function.

ITS2 sequences were processed using the DADA2 ITS2 Pipeline Workflow (v1.8) (https://benjjneb.github.io/dada2/ITS_workflow.html). Primer sequences were removed from the reads with cutadapt (Martin, 2011), quality-trimmed (filterAndTrim: minLen = 50, maxEE = 10, 10; for both *P. acuta* and *M. ampliata* samples to minimize read loss), learning error rates, dereplication, read merging and removing chimeric sequences. Taxonomy was assigned using the customized database (ITS2db_trimmed_derep_dada.fasta) from Claar *et al*., (2020) (https://github.com/baumlab/Claar_etal_2020_SciRep/). Sequences were rarefied to 95% sequencing depth using phyloseq (McMurdie and Holmes, 2013).

For both datasets, alpha diversity indices (Chao1, Shannon-Wiener, and Inverse Simpson) were calculated with the rarefied data. Linear mixed models were generated with log-transformed richness data using lme4 (v1.1-26) (Bates *et al*., 2015), with colony effects accounted for in the models. Post-hoc Tukey HSD tests with emmeans (v1.6.0) (Lenth, 2023) was used to determine the contrasting macroalgal contact type groups contributing to significant changes observed during the calculations. Relative abundances were calculated using the microbiome package (Lahti *et al*., 2018; Neave *et al*., 2017). Beta diversity was assessed using Permutational Multivariate Analysis of Variance (PERMANOVA) and Permutational Analysis of Multivariate Dispersions (PERMDISP) on square-root transformed data using vegan (v2.5-7) (Oksanen *et al*., 2007), with post-hoc pairwise PERMANOVA performed using pairwiseAdonis (v0.4) (Martinez Arbizu, 2017). The strata function in adonis2 was used to account for possible colony effects during calculations. Bray-Curtis-based non-metric multidimensional scaling (nMDS) plots were generated with ggplot2 (v3.3.3) (Wickham, 2016). Differentially abundant bacterial ASVs between the control (no contact) and treatments (direct and water-mediated contact) were identified using DESeq2 (v1.30.0) (Love *et al*., 2014). ITS2 variable region sequences were additionally analysed using the SymPortal analytical framework (v0.3.24) operating on the remote server architecture (SymPortal 2.0; https://symportal.org/) (Hume *et al*., 2019) to further characterise the Symbiodiniaceae communities in the coral samples.

## Acknowledgements

This project is supported by the National Research Foundation, Singapore, and the National Parks Board (NParks), Singapore under its Marine Climate Change Science Programme (MCCS Award NRF-MCCS21-1-5-0001). Any opinions, findings and conclusions or recommendations expressed in this material are those of the author(s) and do not reflect the views of National Research Foundation, Singapore and National Parks Board, Singapore. The Singapore Centre for Environmental Life Sciences Engineering (SCELSE) is funded by the Ministry of Education, Singapore, the National Research Foundation of Singapore, Nanyang Technological University Singapore (NTU) and National University of Singapore (NUS). The authors would like to acknowledge St. John’s Island National Marine Laboratory (SJINML) for providing the facility necessary for conducting the research. The Laboratory is a National Research Infrastructure under the National Research Foundation Singapore. We would also like to thank the SJI Coral team at the NUS Tropical Marine Science Institute (TMSI) for collecting the coral fragments and their experimental support.

## Notes

### Competing Interest Statement

The authors have declared no competing interest.

