## Supplementary Tables for "Contact- and diffusion-based allelopathic interaction of a seaweed with coral holobionts"

**Table S1. Summary of linear mixed models analyses on alpha-diversity indices for *P. acuta* and *M. ampliata* microbiomes based on macroalgal contact type.**

| **Species** | **Location** | **Alpha diversity index** | **D_f_** | **Sum Sq** | **Mean sq** | ***F* value** | ***P_r_* (>*F*)** |
| --- | --- | --- | --- | --- | --- | --- | --- |
| *P. acuta* | Pulau Satumu | Chao1 | 4 | 8.43 | 2.11 | 13.5 | **6.47e-05** |
|  |  | Shannon-Wiener | 4 | 0.841 | 0.210 | 3.74 | **0.027** |
|  |  | Inverse Simpson | 4 | 11.9 | 2.97 | 3.69 | **0.028** |
|  | Kusu Island | Chao1 | 4 | 9.37 | 2.34 | 22.2 | **2.28e-06** |
|  |  | Shannon-Wiener | 4 | 1.40 | 0.351 | 7.73 | **0.0012** |
|  |  | Inverse Simpson | 4 | 11.8 | 2.95 | 5.79 | **0.0045** |
| *M. ampliata* | Pulau Satumu | Chao1 | 4 | 2.46 | 0.614 | 2.14 | 0.12 |
|  |  | Shannon-Wiener | 4 | 0.390 | 0.0975 | 1.10 | 0.39 |
|  |  | Inverse Simpson | 4 | 1.20 | 0.300 | 0.525 | 0.72 |
|  | Kusu Island | Chao1 | 4 | 9.95 | 2.49 | 11.2 | **0.000157** |
|  |  | Shannon-Wiener | 4 | 1.35 | 0.339 | 12.9 | **6.93e-05** |
|  |  | Inverse Simpson | 4 | 16.3 | 4.07 | 16.4 | **1.65e-05** |

* Bold values indicate significant differences when p < 0.05.

**Table S2. Summary of post-hoc Tukey HSD analyses for *P. acuta* and *M. ampliata* microbiomes based on macroalgal contact type.**

| **Species** | **Location** | **Alpha diversity index** | **Pairwise comparison group** | ***P_r_* (>*F*)** |
| --- | --- | --- | --- | --- |
| *P. acuta* | Pulau Satumu | Chao1 | Water-mediated / non-treated control | 0.76 |
|  |  |  | Water-mediated / direct contact | 0.83 |
|  |  |  | Water-mediated / field sample | **0.019** |
|  |  |  | Water-mediated / before contact treatment | **0.0001** |
|  |  |  | Non-treated control / direct contact | 1.00 |
|  |  |  | Non-treated control / field sample | 0.15 |
|  |  |  | Non-treated control / before contact treatment | **0.0008** |
|  |  |  | Direct contact / field sample | 0.12 |
|  |  |  | Direct contact / before contact treatment | **0.0006** |
|  |  |  | Field sample / before contact treatment | 0.16 |
|  |  | Shannon-Wiener | Water-mediated / non-treated control | 1.00 |
|  |  |  | Water-mediated / direct contact | 0.96 |
|  |  |  | Water-mediated / field sample | 0.80 |
|  |  |  | Water-mediated / before contact treatment | 0.075 |
|  |  |  | Non-treated control / direct contact | 0.85 |
|  |  |  | Non-treated control / field sample | 0.93 |
|  |  |  | Non-treated control / before contact treatment | 0.13 |
|  |  |  | Direct contact / field sample | 0.45 |
|  |  |  | Direct contact / before contact treatment | **0.021** |
|  |  |  | Field sample / before contact treatment | 0.54 |
|  |  | Inverse Simpson | Water-mediated / non-treated control | 0.72 |
|  |  |  | Water-mediated / direct contact | 0.95 |
|  |  |  | Water-mediated / field sample | 0.54 |
|  |  |  | Water-mediated / before contact treatment | 0.10 |
|  |  |  | Non-treated control / direct contact | 0.31 |
|  |  |  | Non-treated control / field sample | 0.99 |
|  |  |  | Non-treated control / before contact treatment | 0.63 |
|  |  |  | Direct contact / field sample | 0.21 |
|  |  |  | Direct contact / before contact treatment | **0.027** |
|  |  |  | Field sample / before contact treatment | 0.88 |
|  | Kusu Island | Chao1 | Water-mediated / non-treated control | 1.00 |
|  |  |  | Water-mediated / direct contact | 0.80 |
|  |  |  | Water-mediated / field sample | **< .0001** |
|  |  |  | Water-mediated / before contact treatment | **0.0006** |
|  |  |  | Non-treated control / direct contact | 0.62 |
|  |  |  | Non-treated control / field sample | **< .0001** |
|  |  |  | Non-treated control / before contact treatment | **0.0003** |
|  |  |  | Direct contact / field sample | **0.0002** |
|  |  |  | Direct contact / before contact treatment | **0.0051** |
|  |  |  | Field sample / before contact treatment | 0.46 |
|  |  | Shannon-Wiener | Water-mediated / non-treated control | 1.00 |
|  |  |  | Water-mediated / direct contact | 1.00 |
|  |  |  | Water-mediated / field sample | **0.0087** |
|  |  |  | Water-mediated / before contact treatment | **0.039** |
|  |  |  | Non-treated control / direct contact | 0.98 |
|  |  |  | Non-treated control / field sample | **0.0057** |
|  |  |  | Non-treated control / before contact treatment | **0.026** |
|  |  |  | Direct contact / field sample | **0.016** |
|  |  |  | Direct contact / before contact treatment | 0.070 |
|  |  |  | Field sample / before contact treatment | 0.94 |
|  |  | Inverse Simpson | Water-mediated / non-treated control | 0.72 |
|  |  |  | Water-mediated / direct contact | 0.95 |
|  |  |  | Water-mediated / field sample | 0.54 |
|  |  |  | Water-mediated / before contact treatment | 0.10 |
|  |  |  | Non-treated control / direct contact | 0.31 |
|  |  |  | Non-treated control / field sample | 0.99 |
|  |  |  | Non-treated control / before contact treatment | 0.63 |
|  |  |  | Direct contact / field sample | 0.21 |
|  |  |  | Direct contact / before contact treatment | **0.027** |
|  |  |  | Field sample / before contact treatment | 0.88 |
| *M. ampliata* | Kusu Island | Chao1 | Water-mediated / non-treated control | 0.98 |
|  |  |  | Water-mediated / direct contact | 1.00 |
|  |  |  | Water-mediated / field sample | **0.0037** |
|  |  |  | Water-mediated / before contact treatment | **0.0030** |
|  |  |  | Non-treated control / direct contact | 0.95 |
|  |  |  | Non-treated control / field sample | **0.011** |
|  |  |  | Non-treated control / before contact treatment | **0.009** |
|  |  |  | Direct contact / field sample | **0.0027** |
|  |  |  | Direct contact / before contact treatment | **0.0022** |
|  |  |  | Field sample / before contact treatment | 1.00 |
|  |  | Shannon-Wiener | Water-mediated / non-treated control | 0.89 |
|  |  |  | Water-mediated / direct contact | 1.00 |
|  |  |  | Water-mediated / field sample | 0.063 |
|  |  |  | Water-mediated / before contact treatment | **0.0001** |
|  |  |  | Non-treated control / direct contact | 0.97 |
|  |  |  | Non-treated control / field sample | 0.29 |
|  |  |  | Non-treated control / before contact treatment | **0.0007** |
|  |  |  | Direct contact / field sample | 0.11 |
|  |  |  | Direct contact / before contact treatment | **0.0002** |
|  |  |  | Field sample / before contact treatment | **0.041** |
|  |  | Inverse Simpson | Water-mediated / non-treated control | 0.23 |
|  |  |  | Water-mediated / direct contact | 0.97 |
|  |  |  | Water-mediated / field sample | **0.048** |
|  |  |  | Water-mediated / before contact treatment | **< .0001** |
|  |  |  | Non-treated control / direct contact | 0.51 |
|  |  |  | Non-treated control / field sample | 0.90 |
|  |  |  | Non-treated control / before contact treatment | **0.001** |
|  |  |  | Direct contact / field sample | 0.14 |
|  |  |  | Direct contact / before contact treatment | **< .0001** |
|  |  |  | Field sample / before contact treatment | **0.0058** |

* Bold values indicate significant differences when p < 0.05.

**Table S3. Summary of the *DESeq2* analyses for *P. acuta* microbiomes, showing some of the ASV changes between both control and treated microbiomes.** Green font indicates increase in relative abundance; blue indicates decrease in relative abundance. Shared ASVs between both comparison treatment pairs are in bold.

| **Location** | **Comparison Pair** | **ASV** | **Log2FoldChange** | **Phylum** | **Lowest Taxonomical Classification** |
| --- | --- | --- | --- | --- | --- |
| Pulau Satumu | Control - Direct | **268** | **31.926** | **Proteobacteria** | **Family Alteromonadaceae** |
|  |  | **16** | **30.717** | **Actinobacteriota** | **Family Actinomarinaceae, *Candidatus_Actinomarina sp.*** |
|  |  | **321** | **29.785** | **Desulfobacterota** | ***Desulfobacter sp.*** |
|  |  | 190 | 24.893 | Desulfobacterota | *Desulfobacter sp.* |
|  |  | 1127 | 23.0964 | Verrucomicrobiota | *Lentimonas sp.* |
|  |  | 910 | 22.923 | Proteobacteria | *Sneathiella sp.* |
|  |  | 323 | 22.890 | Planctomycetota | Family Phycisphaeraceae*, SM1A02 sp.* |
|  |  | 882 | 22.286 | Bacteroidota | *Aquibacter sp.* |
|  |  | 900 | 22.260 | Bacteroidota | *Zeaxanthinibacter sp.* |
|  |  | 730 | 22.00853 | Proteobacteria | *Erythrobacter sp.* |
|  |  | 1103 | 21.965 | Bacteroidota | Family Flavobacteriaceae |
|  |  | 824 | 21.794 | Proteobacteria | *Woeseia sp.* |
|  |  | 296 | 21.535 | Proteobacteria | Family Rhodobacteraceae |
|  |  | 352 | 21.412 | Fusobacteriota | *Ilyobacter sp.* |
|  |  | 1139 | 20.929 | Bacteroidota | *Carboxylicivirga sediminis* |
|  |  | 1182 | 20.314 | Planctomycetota | *Pirellula sp.* |
|  |  | 455 | 20.0306 | Proteobacteria | *Dasania sp.* |
|  |  | **647** | **-18.0553** | **Proteobacteria** | **Order Gammaproteobacteria_Incertae_Sedis** |
|  |  | **328** | **-19.216** | **Proteobacteria** | ***Oleiphilus sp.*** |
|  |  | **272** | **-19.295** | **Bacteroidota** | ***Tenacibaculum litopenaei*** |
|  |  | **875** | **-20.207** | **Bacteroidota** | **Family Saprospiraceae** |
|  |  | **416** | **-20.784** | **Proteobacteria** | **Family Rickettsiaceae, *Candidatus_Megaira sp.*** |
|  |  | **1494** | **-20.859** | **Planctomycetota** | **Family Pirellulaceae** |
|  |  | **1927** | **-20.995** | **Proteobacteria** | **Family Porticoccaceae** |
|  |  | **359** | **-21.638** | **Proteobacteria** | ***Kordiimonas sp.*** |
|  |  | **935** | **-21.824** | **Bacteroidota** | ***Algibacter sp.*** |
|  |  | **39** | **-22.0548** | **Bacteroidota** | ***Muricauda sp.*** |
|  |  | **1257** | **-22.138** | **Bacteroidota** | ***Muricauda sp.*** |
|  |  | **386** | **-22.308** | **Proteobacteria** | **Family Diplorickettsiaceae** |
|  |  | **707** | **-22.525** | **Proteobacteria** | ***Woeseia sp.*** |
|  |  | **1330** | **-22.673** | **Bacteroidota** | **Family Flavobacteriaceae** |
|  |  | **173** | **-23.196** | **Verrucomicrobiota** | ***Lentimonas sp.*** |
|  |  | **465** | **-23.333** | **Proteobacteria** | ***Maricurvus nonylphenolicus*** |
|  |  | **667** | **-24.743** | **Verrucomicrobiota** | ***Lentimonas sp.*** |
|  |  | 108 | -25.508 | Proteobacteria | Family Alteromonadaceae |
|  |  | 850 | -26.400 | Proteobacteria | *Pseudomaricurvus alkylphenolicus* |
|  |  | 263 | -28.226 | Proteobacteria | *Kordiimonas sp.* |
|  |  | 29 | -29.367 | Proteobacteria | *Marinobacterium sp.* |
|  |  | 929 | -39.788 | Bacteroidota | *Pedobacter sp.* |
|  |  | 145 | -40.623 | Actinobacteriota | *Micrococcus sp.* |
|  | Control – 5 cm (water-mediated) | **321** | **25.0200** | **Desulfobacterota** | ***Desulfobacter sp.*** |
|  |  | 504 | 22.0556 | Proteobacteria | Family Rhodobacteraceae |
|  |  | 1109 | 21.642 | Bacteroidota | *Reichenbachiella sp.* |
|  |  | **268** | **20.609** | **Proteobacteria** | **Family Alteromonadaceae** |
|  |  | 908 | 20.449 | Proteobacteria | Family Halieaceae |
|  |  | 554 | 20.187 | Proteobacteria | *Altererythrobacter sp.* |
|  |  | 888 | 20.120 | Proteobacteria | Family Cellvibrionaceae, *Candidatus_Endobugula sp.* |
|  |  | 102 | 19.961 | Proteobacteria | *Neptuniibacter marinus* |
|  |  | 215 | 19.721 | Proteobacteria | *Thalassococcus halodurans* |
|  |  | 69 | 19.626 | Cyanobacteria | Family Cyanobiaceae, *Synechococcus_CC9902 sp.* |
|  |  | 1668 | 19.365 | Proteobacteria | *Perspicuibacter marinus* |
|  |  | **16** | **19.172** | **Actinobacteriota** | **Family Actinomarinaceae, *Candidatus_Actinomarina sp.*** |
|  |  | 462 | -19.190 | Chloroflexi | Family Anaerolineaceae |
|  |  | **39** | **-21.280** | **Bacteroidota** | ***Muricauda sp.*** |
|  |  | 1067 | -21.299 | Proteobacteria | Family Thiohalorhabdaceae |
|  |  | 427 | -21.342 | Proteobacteria | *Oleiphilus sp.* |
|  |  | **465** | **-21.560** | **Proteobacteria** | ***Maricurvus nonylphenolicus*** |
|  |  | **667** | **-21.613** | **Verrucomicrobiota** | ***Lentimonas sp.*** |
|  |  | **359** | **-21.654** | **Proteobacteria** | ***Kordiimonas sp.*** |
|  |  | **707** | **-21.859** | **Proteobacteria** | ***Woeseia sp.*** |
|  |  | **1257** | **-21.983** | **Bacteroidota** | ***Muricauda sp.*** |
|  |  | **1330** | **-22.0517** | **Bacteroidota** | **Family Flavobacteriaceae** |
|  |  | 897 | -22.135 | Proteobacteria | Order Gammaproteobacteria_Incertae_Sedis |
|  |  | **173** | **-22.390** | **Verrucomicrobiota** | ***Lentimonas sp.*** |
|  |  | **1494** | **-22.555** | **Planctomycetota** | **Family Pirellulaceae** |
|  |  | **1927** | **-22.582** | **Proteobacteria** | **Family Porticoccaceae** |
|  |  | **935** | **-22.664** | **Bacteroidota** | ***Algibacter sp.*** |
|  |  | **875** | **-22.675** | **Bacteroidota** | **Family Saprospiraceae** |
|  |  | **647** | **-22.881** | **Proteobacteria** | **Order Gammaproteobacteria_Incertae_Sedis** |
|  |  | 219 | -23.187 | Proteobacteria | Family Alteromonadaceae |
|  |  | 1015 | -23.193 | Proteobacteria | *Kordiimonas sp.* |
|  |  | 1357 | -23.268 | Proteobacteria | *Erythrobacter sp.* |
|  |  | 26 | -23.478 | Cyanobacteria | Family Cyanobiaceae, *Synechococcus_CC9902 sp.* |
|  |  | 327 | -23.884 | Verrucomicrobiota | *Rubritalea sp.* |
|  |  | **386** | **-24.882** | **Proteobacteria** | **Family Diplorickettsiaceae** |
|  |  | **272** | **-25.0859** | **Bacteroidota** | ***Tenacibaculum litopenaei*** |
|  |  | **416** | **-25.435** | **Proteobacteria** | **Family Rickettsiaceae, *Candidatus_Megaira sp.*** |
|  |  | **328** | **-25.672** | **Proteobacteria** | ***Oleiphilus sp.*** |
| Kusu Island | Control - Direct | **16** | **37.505** | **Actinobacteriota** | **Family Actinomarinaceae, *Candidatus_Actinomarina sp.*** |
|  |  | **201** | **29.418** | **Proteobacteria** | **Family Hyphomonadaceae** |
|  |  | **772** | **28.273** | **Proteobacteria** | **Order Parvibaculales, Family PS1_clade** |
|  |  | **755** | **26.773** | **Proteobacteria** | ***Pseudomonas sp.*** |
|  |  | **388** | **26.373** | **Proteobacteria** | ***Arenicella sp.*** |
|  |  | **264** | **26.279** | **Proteobacteria** | ***Litoribrevibacter sp*** |
|  |  | 350 | 26.228 | Verrucomicrobiota | *Lentimonas sp.* |
|  |  | **2091** | **25.812** | **Proteobacteria** | ***Pseudahrensia sp.*** |
|  |  | 173 | 25.587 | Verrucomicrobiota | *Lentimonas sp.* |
|  |  | **3890** | **25.423** | **Proteobacteria** | ***Marinicella sp.*** |
|  |  | **583** | **24.802** | **Proteobacteria** | **Family Rhodobacteraceae** |
|  |  | 343 | 23.869 | Proteobacteria | *Aestuariibacter sp.* |
|  |  | 1904 | 23.363 | Desulfobacterota | *Desulfobacter sp.* |
|  |  | 396 | 23.156 | Proteobacteria | Family Cellvibrionaceae, *Candidatus_Endobugula sp.* |
|  |  | 340 | 23.127 | Campilobacterota | Family Arcobacteraceae |
|  |  | 131 | 22.916 | Proteobacteria | *Pseudohongiella sp.* |
|  |  | 884 | 22.822 | Verrucomicrobiota | Order Verrucomicrobiales, Family DEV007 |
|  |  | 311 | 22.775 | Proteobacteria | Family Porticoccaceae. *C1-B045 sp.* |
|  |  | 93 | 22.683 | Proteobacteria | *Marichromatium sp.* |
|  |  | 660 | 22.652 | Proteobacteria | *Thalassotalea sp.* |
|  |  | 626 | 22.599 | Proteobacteria | *Kordiimonas sp.* |
|  |  | 351 | 22.109 | Proteobacteria | Order Cellvibrionales, Family BD2-7 |
|  |  | 47 | 22.0631 | Cyanobacteria | Family Cyanobiaceae, *Cyanobium_PCC-6307 sp.* |
|  |  | 296 | 21.642 | Proteobacteria | Family Rhodobacteraceae |
|  |  | 611 | 21.532 | Verrucomicrobiota | *Lentimonas sp.* |
|  |  | 1210 | 21.450 | Bacteroidota | Family Cryomorphaceae |
|  |  | 920 | 20.695 | Bacteroidota | *Robiginitalea sp.* |
|  |  | 435 | 20.653 | Proteobacteria | *Motiliproteus sp.* |
|  |  | 880 | 19.402 | Planctomycetota | *Rhodopirellula sp.* |
|  |  | 153 | 19.0595 | Proteobacteria | *Kordiimonas sp.* |
|  |  | **1006** | **-17.600** | **Verrucomicrobiota** | ***Rubritalea sp.*** |
|  |  | **441** | **-18.130** | **Planctomycetota** | **Family Phycisphaeraceae, *SM1A02 sp.*** |
|  |  | **606** | **-19.618** | **Proteobacteria** | ***Arenicella sp.*** |
|  |  | **178** | **-20.165** | **Proteobacteria** | **Family Vibrionaceae** |
|  |  | **1205** | **-20.696** | **Bacteroidota** | ***Crocinitomix sp.*** |
|  |  | **416** | **-21.00132** | **Proteobacteria** | **Family Rickettsiaceae, *Candidatus_Megaira sp.*** |
|  |  | **934** | **-21.0398** | **Dependentiae** | **Family Vermiphilaceae** |
|  |  | **2078** | **-21.107** | **Proteobacteria** | **Family Cellvibrionaceae, *Candidatus_Endobugula sp.*** |
|  |  | **289** | **-21.405** | **Proteobacteria** | ***Thalassotalea sp.*** |
|  |  | **428** | **-22.394** | **Bacteroidota** | ***Muricauda sp.*** |
|  |  | **923** | **-22.718** | **Proteobacteria** | **Family Cellvibrionaceae, *Candidatus_Endobugula sp.*** |
|  |  | **455** | **-22.796** | **Proteobacteria** | ***Dasania sp.*** |
|  |  | **756** | **-22.805** | **Proteobacteria** | **Family Piscirickettsiaceae. *Candidatus_Endoecteinascidia sp.*** |
|  |  | **1037** | **-22.854** | **Proteobacteria** | ***Ferrimonas kyonanensis*** |
|  |  | **386** | **-23.541** | **Proteobacteria** | **Family Diplorickettsiaceae** |
|  |  | 1527 | -25.226 | Proteobacteria | Family Rhodobacteraceae |
|  |  | 2475 | -25.535 | Proteobacteria | Family Halieaceae |
|  |  | 200 | -28.685 | Bacteroidota | Family Flavobacteriaceae, *NS4_marine_group sp.* |
|  |  | 1631 | -28.686 | Proteobacteria | Order Gammaproteobacteria_Incertae_Sedis |
|  |  | 357 | -28.697 | Proteobacteria | *Ralstonia sp.* |
|  | Control – 5 cm (water-mediated) | 645 | 23.558 | Verrucomicrobiota | *Lentimonas sp.* |
|  |  | **2091** | **22.613** | **Proteobacteria** | ***Pseudahrensia sp.*** |
|  |  | **583** | **22.346** | **Proteobacteria** | **Family Rhodobacteraceae** |
|  |  | **388** | **22.260** | **Proteobacteria** | ***Arenicella sp.*** |
|  |  | 494 | 22.259 | Verrucomicrobiota | *Lentimonas sp.* |
|  |  | 710 | 22.129 | Proteobacteria | *Erythrobacter sp.* |
|  |  | 577 | 22.0454 | Proteobacteria | Family Rhodobacteraceae |
|  |  | **3890** | **21.818** | **Proteobacteria** | ***Marinicella sp.*** |
|  |  | 256 | 21.798 | Proteobacteria | Family Terasakiellaceae |
|  |  | 865 | 21.508 | Proteobacteria | *Ruegeria sp.* |
|  |  | 1002 | 21.297 | Bacteroidota | *Tenacibaculum sp.* |
|  |  | 902 | 21.183 | Proteobacteria | Family Cellvibrionaceae, *Candidatus_Endobugula sp.* |
|  |  | **201** | **20.851** | **Proteobacteria** | **Family Hyphomonadaceae** |
|  |  | 1409 | 20.525 | Planctomycetota | Family Phycisphaeraceae, *Urania-1B-19_marine_sediment_group sp.* |
|  |  | 212 | 20.139 | Proteobacteria | Family Rhodobacteraceae |
|  |  | 522 | 19.874 | Proteobacteria | *Woeseia sp.* |
|  |  | 1480 | 19.284 | Bacteroidota | *Portibacter sp.* |
|  |  | **772** | **19.0432** | **Proteobacteria** | **Order Parvibaculales, Family PS1_clade** |
|  |  | **16** | **19.00193** | **Actinobacteriota** | **Family Actinomarinaceae, *Candidatus_Actinomarina sp.*** |
|  |  | 1062 | 17.872 | Planctomycetota | *Rubripirellula sp.* |
|  |  | **755** | **17.271** | **Proteobacteria** | ***Pseudomonas sp.*** |
|  |  | 108 | 17.0567 | Proteobacteria | Family Alteromonadaceae |
|  |  | **264** | **17.00604** | **Proteobacteria** | ***Litoribrevibacter sp.*** |
|  |  | 507 | 15.906 | Proteobacteria | *Sphingomonas sp.* |
|  |  | **1037** | **-17.286** | **Proteobacteria** | ***Ferrimonas kyonanensis*** |
|  |  | **923** | **-17.909** | **Proteobacteria** | **Family Cellvibrionaceae, *Candidatus_Endobugula sp.*** |
|  |  | 23 | -19.510 | Proteobacteria | Order SAR11_clade, Family Clade_I, *Clade_Ib sp.* |
|  |  | **756** | **-20.140** | **Proteobacteria** | **Family Piscirickettsiaceae, *Candidatus_Endoecteinascidia sp.*** |
|  |  | **428** | **-20.169** | **Bacteroidota** | ***Muricauda sp.*** |
|  |  | **455** | **-20.481** | **Proteobacteria** | ***Dasani sp.*** |
|  |  | **416** | **-20.651** | **Proteobacteria** | **Family Rickettsiaceae, *Candidatus_Megaira sp.*** |
|  |  | 504 | -21.331 | Proteobacteria | Family Rhodobacteraceae |
|  |  | **386** | **-21.808** | **Proteobacteria** | **Family Diplorickettsiaceae** |
|  |  | **441** | **-21.843** | **Planctomycetota** | **Family Phycisphaeraceae, *SM1A02 sp.*** |
|  |  | 1526 | -22.192 | Bdellovibrionota | Family Bdellovibrionaceae, *OM27_clade sp.* |
|  |  | **2078** | **-22.248** | **Proteobacteria** | **Family Cellvibrionaceae, *Candidatus_Endobugula sp.*** |
|  |  | **289** | **-22.462** | **Proteobacteria** | ***Thalassotalea sp.*** |
|  |  | 312 | -22.477 | Proteobacteria | *Kiloniella sp.* |
|  |  | 2523 | -22.504 | Myxococcota | Family Nannocystaceae |
|  |  | **1205** | **-22.660** | **Bacteroidota** | ***Crocinitomix sp.*** |
|  |  | **934** | **-23.464** | **Dependentiae** | **Family Vermiphilaceae** |
|  |  | **1006** | **-23.500** | **Verrucomicrobiota** | ***Rubritalea sp.*** |
|  |  | **606** | **-23.813** | **Proteobacteria** | ***Arenicella sp.*** |
|  |  | 1094 | -24.135 | Proteobacteria | *Aestuariibacter sp.* |
|  |  | **178** | **-24.518** | **Proteobacteria** | **Family Vibrionaceae** |
|  |  | 432 | -24.728 | Desulfobacterota | *Desulfofrigus sp.* |

**Table S4. Summary of the *DESeq2* analyses for *M. ampliata* microbiomes, showing some of the ASV changes between both control and treated microbiomes.** Green font indicates increase in relative abundance; blue indicates decrease in relative abundance. Shared ASVs between both comparison treatment pairs are in bold.

| **Location** | **Comparison Pair** | **ASV** | **Log2FoldChange** | **Phylum** | **Lowest Taxonomical Classification** |
| --- | --- | --- | --- | --- | --- |
| Pulau Satumu | Control - Direct | **83** | **30.876** | **Proteobacteria** | ***Pseudoalteromonas sp.*** |
|  |  | **2053** | **29.561** | **Verrucomicrobiota** | **Family DEV007** |
|  |  | **465** | **26.810** | **Proteobacteria** | ***Maricurvus nonylphenolicus*** |
|  |  | **1718** | **26.529** | **Proteobacteria** | ***Arenicella sp.*** |
|  |  | 1458 | 23.650 | Bacteroidota | *Crocinitomix sp.* |
|  |  | 1228 | 23.0246 | Bdellovibrionota | Family Bdellovibrionaceae, *OM27_clade sp.* |
|  |  | 1283 | 22.987 | Bacteroidota | *Muricauda sp.* |
|  |  | 36 | 22.947 | Proteobacteria | Family Vibrionaceae |
|  |  | 264 | 22.751 | Proteobacteria | *Litoribrevibacter sp.* |
|  |  | 475 | 22.434 | Proteobacteria | *Shimia sp.* |
|  |  | 858 | 22.365 | Proteobacteria | *Aliikangiella sp.* |
|  |  | 343 | 22.356 | Proteobacteria | *Aestuariibacter sp.* |
|  |  | 405 | 22.0864 | Proteobacteria | Family Cellvibrionaceae |
|  |  | 884 | 21.859 | Verrucomicrobiota | Family DEV007 |
|  |  | 504 | 21.421 | Proteobacteria | Family Rhodobacteraceae |
|  |  | 865 | 21.394 | Proteobacteria | *Ruegeria sp.* |
|  |  | 394 | 20.869 | Proteobacteria | *Aestuariicella sp.* |
|  |  | 539 | 20.239 | Planctomycetota | *Blastopirellula sp.* |
|  |  | 507 | 18.633 | Proteobacteria | *Sphingomonas sp.* |
|  |  | **11** | **-18.789** | **Proteobacteria** | **Order SAR11_clade, Family Clade_I, *Clade_Ia sp.*** |
|  |  | **767** | **-20.871** | **Proteobacteria** | **Family Cellvibrionaceae, *Candidatus_Endobugula sp.*** |
|  |  | **921** | **-21.0794** | **Planctomycetota** | ***Blastopirellula sp.*** |
|  |  | **258** | **-22.269** | **Proteobacteria** | ***Woeseia sp.*** |
|  |  | **368** | **-22.444** | **Proteobacteria** | ***Arenicella sp.*** |
|  |  | **233** | **-22.721** | **Proteobacteria** | ***Ruegeria conchae*** |
|  |  | **212** | **-22.800** | **Proteobacteria** | **Family Rhodobacteraceae** |
|  |  | 647 | -23.262 | Proteobacteria | Order Gammaproteobacteria_Incertae_Sedis |
|  |  | **136** | **-25.193** | **Proteobacteria** | **Family Rhodobacteraceae** |
|  |  | **254** | **-26.501** | **Cyanobacteria** | ***Trichodesmium_IMS101 sp.*** |
|  |  | 412 | -26.679 | Planctomycetota | *Phycisphaera sp.* |
|  |  | 749 | -26.694 | Proteobacteria | Family Nitrosococcaceae, *AqS1 sp.* |
|  |  | 1973 | -26.796 | Proteobacteria | Family Alteromonadaceae |
|  |  | 923 | -26.952 | Proteobacteria | Family Cellvibrionaceae, *Candidatus_Endobugula sp.* |
|  |  | 893 | -28.741 | Bacteroidota | *Maritimimonas sp.* |
|  |  | 8 | -29.733 | Actinobacteriota | Family Actinomarinaceae, *Candidatus_Actinomarina sp.* |
|  |  | 1244 | -30.307 | Proteobacteria | Family Spongiibacteraceae*, BD1-7_clade sp.* |
|  |  | 506 | -35.561 | Proteobacteria | *Halioglobus sp.* |
|  | Control – 5 cm (water-mediated) | 150 | 23.364 | Verrucomicrobiota | *Lentimonas sp.* |
|  |  | **465** | **23.288** | **Proteobacteria** | ***Maricurvus nonylphenolicus*** |
|  |  | **1718** | **22.327** | **Proteobacteria** | ***Arenicella sp.*** |
|  |  | 674 | 21.959 | Proteobacteria | Family Rhizobiaceae |
|  |  | **2053** | **21.593** | **Verrucomicrobiota** | **Family DEV007** |
|  |  | **83** | **21.0760** | **Proteobacteria** | ***Pseudoalteromonas sp.*** |
|  |  | 824 | 20.629 | Proteobacteria | *Woeseia sp.* |
|  |  | 620 | 20.508 | Proteobacteria | Family Rhodobacteraceae |
|  |  | 614 | 20.370 | Proteobacteria | Family Cellvibrionaceae, *Candidatus_Endobugula sp.* |
|  |  | 596 | 20.319 | Proteobacteria | *Aliikangiella sp.* |
|  |  | 367 | 20.249 | Proteobacteria | *Halioglobus sp.* |
|  |  | 850 | 19.995 | Proteobacteria | *Pseudomaricurvus alkylphenolicus* |
|  |  | 69 | 19.665 | Cyanobacteria | Family Cyanobiaceae, *Synechococcus_CC9902 sp.* |
|  |  | 444 | 19.482 | Proteobacteria | Order Gammaproteobacteria_Incertae_Sedis |
|  |  | 522 | 19.456 | Proteobacteria | *Woeseia sp.* |
|  |  | 882 | 19.244 | Bacteroidota | *Aquibacter sp.* |
|  |  | 880 | 18.842 | Planctomycetota | *Rhodopirellula sp.* |
|  |  | 908 | 18.116 | Proteobacteria | Family Halieaceae |
|  |  | 804 | 17.548 | Proteobacteria | *Labrenzia marina* |
|  |  | 822 | -19.117 | Verrucomicrobiota | Family DEV007 |
|  |  | 617 | -19.244 | Proteobacteria | *Algicola bacteriolytica* |
|  |  | 145 | -20.391 | Actinobacteriota | *Micrococcus sp.* |
|  |  | **11** | **-20.436** | **Proteobacteria** | **Order SAR11_clade, Family Clade_I, *Clade_Ia sp.*** |
|  |  | **368** | **-20.835** | **Proteobacteria** | ***Arenicella sp.*** |
|  |  | **136** | **-20.976** | **Proteobacteria** | **Family Rhodobacteraceae** |
|  |  | **258** | **-21.356** | **Proteobacteria** | ***Woeseia sp.*** |
|  |  | 818 | -21.725 | Bacteroidota | Family Cyclobacteriaceae |
|  |  | **233** | **-21.823** | **Proteobacteria** | ***Ruegeria conchae*** |
|  |  | 352 | -22.0667 | Fusobacteriota | *Ilyobacter sp.* |
|  |  | 1954 | -22.0679 | Verrucomicrobiota | Family DEV007 |
|  |  | **921** | **-22.144** | **Planctomycetota** | ***Blastopirellula sp.*** |
|  |  | 764 | -22.780 | Myxococcota | *Family Nannocystaceae* |
|  |  | **767** | **-22.877** | **Proteobacteria** | **Family Cellvibrionaceae, *Candidatus_Endobugula sp.*** |
|  |  | 1545 | -22.982 | Bacteroidota | *Family Saprospiraceae* |
|  |  | 874 | -23.656 | Proteobacteria | Order Rhizobiales, Family Stappiaceae |
|  |  | 1148 | -24.0717 | Hydrogenedentes | *Family Hydrogenedensaceae* |
|  |  | **212** | **-24.397** | **Proteobacteria** | **Family Rhodobacteraceae** |
|  |  | **254** | **-24.702** | **Cyanobacteria** | **Family Phormidiaceae, *Trichodesmium_IMS101 sp.*** |
|  |  | 345 | -25.108 | Proteobacteria | *Cohaesibacter gelatinilyticus* |
| Kusu Island | Control - Direct | **1272** | **30.758** | **Proteobacteria** | **Order Caulobacterales, Family Hyphomonadaceae** |
|  |  | **328** | **29.188** | **Proteobacteria** | ***Oleiphilus sp.*** |
|  |  | **440** | **26.224** | **Verrucomicrobiota** | ***Lentimonas sp.*** |
|  |  | **647** | **26.0450** | **Proteobacteria** | **Order Gammaproteobacteria_Incertae_Sedis** |
|  |  | **2362** | **25.308** | **Proteobacteria** | **Order Rickettsiales, Family Fokiniaceae*, MD3-55 sp.*** |
|  |  | 703 | 22.794 | Proteobacteria | *Thalassotalea sp.* |
|  |  | 730 | 22.605 | Proteobacteria | *Erythrobacter sp.* |
|  |  | 2215 | 21.979 | Verrucomicrobiota | Family Rubritaleaceae |
|  |  | 1822 | 21.501 | Planctomycetota | Family Phycisphaeraceae*, SM1A02 sp.* |
|  |  | 1876 | 21.340 | Planctomycetota | *Phycisphaera sp.* |
|  |  | 435 | 20.919 | Proteobacteria | *Motiliproteus sp.* |
|  |  | 758 | 20.834 | Proteobacteria | *Leisingera sp.* |
|  |  | 858 | 20.599 | Proteobacteria | *Aliikangiella sp.* |
|  |  | 721 | 20.353 | Proteobacteria | Family Alteromonadaceae |
|  |  | 1196 | 19.840 | Proteobacteria | *Mesorhizobium sp.* |
|  |  | 1169 | 19.231 | Planctomycetota | *Blastopirellula sp.* |
|  |  | **982** | **-19.135** | **Proteobacteria** | ***Kordiimonas sp.*** |
|  |  | **1625** | **-19.811** | **Planctomycetota** | **Family Gimesiaceae** |
|  |  | **33** | **-20.167** | **Proteobacteria** | **Order Rhodospirillales, Family AEGEAN-169_marine_group** |
|  |  | **1915** | **-20.202** | **Myxococcota** | **Family Sandaracinaceae** |
|  |  | **1512** | **-20.804** | **Planctomycetota** | ***Blastopirellula sp.*** |
|  |  | **394** | **-20.883** | **Proteobacteria** | ***Aestuariicella sp.*** |
|  |  | **902** | **-20.961** | **Proteobacteria** | **Family Cellvibrionaceae, *Candidatus_Endobugula sp.*** |
|  |  | **1114** | **-21.212** | **Proteobacteria** | ***Kordiimonas sp.*** |
|  |  | **1908** | **-21.377** | **Verrucomicrobiota** | **Order Chlamydiales, Family Simkaniaceae** |
|  |  | **393** | **-22.141** | **Bdellovibrionota** | ***Peredibacter sp.*** |
|  |  | **1265** | **-23.201** | **Proteobacteria** | ***Oceanimonas sp.*** |
|  |  | **789** | **-23.325** | **Proteobacteria** | ***Neptuniibacter sp.*** |
|  |  | **649** | **-23.434** | **Bacteroidota** | ***Muricauda sp.*** |
|  |  | **344** | **-23.592** | **Proteobacteria** | ***Porticoccus sp.*** |
|  |  | **628** | **-23.638** | **Bacteroidota** | **Family Flavobacteriaceae*, Wenyingzhuangia sp.*** |
|  |  | 1759 | -25.803 | Chloroflexi | Family Anaerolineaceae, *GWD2-49-16 sp.* |
|  |  | 2004 | -26.455 | Acidobacteriota | *Acanthopleuribacter sp.* |
|  |  | 2190 | -26.906 | Proteobacteria | Family Nitrincolaceae |
|  |  | 755 | -27.0440 | Proteobacteria | *Pseudomonas sp.* |
|  |  | 160 | -28.0725 | Proteobacteria | *Halioxenophilus aromaticivorans* |
|  |  | 1337 | -29.763 | Proteobacteria | *Methylophaga sp.* |
|  |  | 2858 | -30.384 | Proteobacteria | Family Methylophagaceae |
|  |  | 69 | -35.103 | Cyanobacteria | Family Cyanobiaceae*, Synechococcus_CC9902 sp.* |
|  | Control – 5 cm (water-mediated) | **328** | **24.778** | **Proteobacteria** | ***Oleiphilus sp.*** |
|  |  | **440** | **23.656** | **Verrucomicrobiota** | ***Lentimonas sp.*** |
|  |  | **2362** | **21.391** | **Proteobacteria** | **Order Rickettsiales, Family Fokiniaceae, *MD3-55 sp.*** |
|  |  | **1272** | **21.246** | **Proteobacteria** | **Order Caulobacterales, Family Hyphomonadaceae** |
|  |  | 2107 | 21.178 | Bacteroidota | *Reichenbachiella sp.* |
|  |  | 1034 | 21.0917 | Proteobacteria | *Altererythrobacter sp.* |
|  |  | 1177 | 21.0859 | Planctomycetota | *Blastopirellula sp.* |
|  |  | 663 | 20.946 | Proteobacteria | Order Cellvibrionales, Family Halieaceae |
|  |  | 752 | 20.826 | Bacteroidota | Family Flavobacteriaceae |
|  |  | 960 | 20.507 | Verrucomicrobiota | *Lentimonas sp.* |
|  |  | 653 | 20.474 | Verrucomicrobiota | *Lentimonas sp.* |
|  |  | 193 | 20.278 | Proteobacteria | *Marinobacterium sp.* |
|  |  | 577 | 20.263 | Proteobacteria | Family Rhodobacteraceae |
|  |  | 1140 | 20.229 | Bacteroidota | *Muricauda sp.* |
|  |  | 795 | 19.873 | Verrucomicrobiota | *Lentimonas sp.* |
|  |  | 11 | 19.677 | Proteobacteria | Order SAR11_clade, Family Clade_I, *Clade_Ia sp.* |
|  |  | 1069 | 19.579 | Verrucomicrobiota | *Lentimonas sp.* |
|  |  | 818 | 19.505 | Bacteroidota | Family Cyclobacteriaceae |
|  |  | 391 | 19.0539 | Proteobacteria | *Sneathiella sp.* |
|  |  | 985 | 17.816 | Proteobacteria | *Woeseia sp.* |
|  |  | **647** | **17.539** | **Proteobacteria** | **Order Gammaproteobacteria_Incertae_Sedis** |
|  |  | 290 | 16.893 | Proteobacteria | Family Cellvibrionaceae, *Halioxenophilus sp.* |
|  |  | 329 | 15.748 | Planctomycetota | Family Pirellulaceae |
|  |  | **1908** | **-17.155** | **Verrucomicrobiota** | **Order Chlamydiales, Family Simkaniaceae** |
|  |  | **1625** | **-20.126** | **Planctomycetota** | **Family Gimesiaceae** |
|  |  | **982** | **-20.909** | **Proteobacteria** | ***Kordiimonas sp.*** |
|  |  | **649** | **-20.959** | **Bacteroidota** | ***Muricauda sp.*** |
|  |  | 2061 | -21.156 | Bacteroidota | Family Flavobacteriaceae |
|  |  | **1265** | **-21.263** | **Proteobacteria** | ***Oceanimonas sp.*** |
|  |  | **393** | **-21.572** | **Bdellovibrionota** | ***Peredibacter sp.*** |
|  |  | **1915** | **-21.695** | **Myxococcota** | **Family Sandaracinaceae** |
|  |  | **628** | **-22.00845** | **Bacteroidota** | **Family Flavobacteriaceae, *Wenyingzhuangia sp.*** |
|  |  | 444 | -22.137 | Proteobacteria | Order Gammaproteobacteria_Incertae_Sedis |
|  |  | **1512** | **-22.163** | **Planctomycetota** | ***Blastopirellula sp.*** |
|  |  | **1114** | **-22.187** | **Proteobacteria** | ***Kordiimonas sp.*** |
|  |  | 2659 | -22.274 | Planctomycetota | Family Pirellulaceae, *Pir4_lineage sp*. |
|  |  | **344** | **-22.400** | **Proteobacteria** | ***Porticoccus sp.*** |
|  |  | 368 | -22.638 | Proteobacteria | *Arenicella sp.* |
|  |  | **902** | **-22.672** | **Proteobacteria** | **Family Cellvibrionaceae, *Candidatus_Endobugula sp.*** |
|  |  | **789** | **-22.687** | **Proteobacteria** | ***Neptuniibacter sp.*** |
|  |  | 3097 | -22.735 | Bacteroidota | Family Flavobacteriaceae |
|  |  | 1533 | -22.763 | Verrucomicrobiota | *Rubritalea sp.* |
|  |  | 212 | -22.775 | Proteobacteria | Family Rhodobacteraceae |
|  |  | 1749 | -22.996 | Cyanobacteria | Family Phormidesmiaceae |
|  |  | **394** | **-23.0132** | **Proteobacteria** | ***Aestuariicella sp.*** |
|  |  | **33** | **-23.472** | **Proteobacteria** | **Order Rhodospirillales, Family AEGEAN-169_marine_group** |
|  |  | 528 | -24.816 | Verrucomicrobiota | *Rubritalea sp.* |
|  |  | 829 | -24.856 | Cyanobacteria | Family Nodosilineaceae*, MBIC10086 sp.* |

**Table S5. Summary of linear mixed models analyses on alpha-diversity indices for *P. acuta* and *M. ampliata* Symbiodiniaceae communities based on macroalgal contact type.**

| **Species** | **Location** | **Alpha diversity index** | **D_f_** | **Sum Sq** | **Mean sq** | ***F* value** | ***P_r_* (>*F*)** |
| --- | --- | --- | --- | --- | --- | --- | --- |
| *P. acuta* | Pulau Satumu | Chao1 | 4 | 1.1146 | 0.2787 | 4.4603 | **0.01414** |
|  |  | Shannon-Wiener | 4 | 1.3262 | 0.3315 | 1.1781 | 0.3597 |
|  |  | Inverse Simpson | 4 | 0.4739 | 0.1185 | 1.0664 | 0.4068 |
|  | Kusu Island | Chao1 | 4 | 1.6933 | 0.4233 | 1.4039 | 0.2774 |
|  |  | Shannon-Wiener | 4 | 0.7107 | 0.1777 | 0.4802 | 0.7499 |
|  |  | Inverse Simpson | 4 | 0.8183 | 0.2046 | 1.9848 | 0.1455 |
| *M. ampliata* | Pulau Satumu | Chao1 | 4 | 0.4085 | 0.1021 | 2.9167 | 0.06725 |
|  |  | Shannon-Wiener | 4 | 0.02457 | 0.006142 | 2.5832 | 0.09088 |
|  |  | Inverse Simpson | 4 | 0.007210 | 0.001803 | 1.5426 | 0.2510 |
|  | Kusu Island | Chao1 | 4 | 0.2679 | 0.06698 | 1.3236 | 0.3090 |
|  |  | Shannon-Wiener | 4 | 0.03572 | 0.008929 | 1.3287 | 0.3068 |
|  |  | Inverse Simpson | 4 | 0.0445 | 0.01113 | 0.6637 | 0.6272 |

* Bold values indicate significant differences when p < 0.05.

**Table S6. Summary of post-hoc Tukey HSD analyses for *P. acuta* Symbiodiniaceae communities based on macroalgal contact type.**

| **Species** | **Location** | **Alpha diversity index** | **Pairwise comparison group** | **Significant? (Y/N)** |
| --- | --- | --- | --- | --- |
| *P. acuta* | Pulau Satumu | Chao1 | Water-mediated / non-treated control | **N** |
|  |  |  | Water-mediated / direct contact | **Y** |
|  |  |  | Water-mediated / field sample | **Y** |
|  |  |  | Water-mediated / before contact treatment | **N** |
|  |  |  | Non-treated control / direct contact | **N** |
|  |  |  | Non-treated control / field sample | **N** |
|  |  |  | Non-treated control / before contact treatment | **N** |
